# Anaerobic fungi in the tortoise alimentary tract illuminate early stages of host-fungal symbiosis and *Neocallimastigomycota* evolution

**DOI:** 10.1101/2023.08.25.554870

**Authors:** Carrie J. Pratt, Casey H. Meili, Adrienne L. Jones, Darian K. Jackson, Emma E. England, Yan Wang, Steve Hartson, Janet Rogers, Mostafa S. Elshahed, Noha H. Youssef

## Abstract

The anaerobic gut fungi (AGF, *Neocallimastigomycota*) reside in the alimentary tract of herbivores. While their presence in mammals is well documented, evidence for their occurrence in non-mammalian hosts is currently sparse. Here we report on AGF communities in tortoises (family *Testudinidae*). Culture-independent surveys of tortoise fecal samples identified a unique AGF community, with three novel deep-branching genera representing >90% of sequences in most samples. Representatives of all genera were successfully isolated under strict anaerobic conditions at 30^°^C or 39^°^C. Transcriptomics-enabled phylogenomic and molecular dating analysis indicated an ancient, deep-branching position in the AGF tree for these genera, with an evolutionary divergence time estimate of 104-112 million years ago (Mya). Such estimates push the establishment of animal- *Neocallimastigomycota* symbiosis from the early Paleogene (67 Mya) to the early Cretaceous (112 Mya). Further, compared to their mammalian counterparts, tortoise-associated isolates exhibited a more limited capacity for plant polysaccharides metabolism and lacked genes encoding several carbohydrate active enzyme (CAZyme) families mediating their degradation. Finally, we demonstrate that the observed curtailed degradation capacities and reduced CAZyme repretoire in tortoise-associated AGF is driven by the paucity of horizontal gene transfer (HGT) in tortoise-associated AGF genomes, compared to the massive HGT occurrence in mammalian AGF taxa. The reduced CAZyome and overall secretory machinery observed is also reflected in an altered cellulosomal production capacity in tortoise-associated AGF. Our findings provide novel insights into the scope of phylogenetic diversity, ecological distribution, evolutionary history, evolution of fungal-host nutritional symbiosis, and dynamics of genes and traits acquisition in *Neocallimastigomycota*.

**Significance:** Anaerobic gut fungi (AGF) are encountered in the rumen and hindgut of mammalian herbivores. However, their occurrence outside their canonical mammalian hosts is currently unclear. We report the identification, isolation, and characterization of novel, deep-branching AGF genera from tortoises. Such discovery expands the phylogenetic diversity and host range of the AGF and revises estimates of the phylum’s evolutionary time to the early Cretaceous (112 Mya). We also demonstrate that tortoise-sourced AGF lack multiple metabolic features compared to their mammalian counterparts, and identify the relative paucity of HGT events in tortoise-associated genera as a major factor underpinning such differences. Our results alter our understanding of the scope of phylogenetic diversity, ecological distribution, and evolutionary history of the AGF.

## Introduction

Microbial communities play a crucial role in the digestive process in herbivores by mediating the breakdown of substrates recalcitrant to their hosts’ enzymes (1–3). The establishment of herbivore-microbiome nutritional symbiosis was associated with the evolution of dedicated digestive chambers e.g. enlarged hindgut, diverticula, and rumen, and longer feed retention times to improve the efficiency of the digestion process (4–7). A complex community of microorganisms in the herbivorous gastrointestinal tract (GIT) breaks down plant biomass to absorbable end products (3). So far, greater emphasis has been placed on the study of bacterial and archaeal members of the community (8–14, 15, 16), compared to microbial eukaryotes (protozoa and fungi). Nevertheless, the role of eukaryotes in the herbivorous gut is increasingly being recognized (17–21).

The anaerobic gut fungi (AGF, *Neocallimastigomycota*) are integral and ubiquitous constituents of the GIT community in mammalian herbivores (22–27). Notably, while chiefly investigated in mammalian hosts, microbiome-enabled herbivory and associated GIT structural features conducive to AGF establishment also occur in multiple non-mammalian herbivores. One of the potential non-mammalian AGF hosts are tortoises, members of the family *Testudinidae*, order *Testudines* (28). Tortoises are terrestrial herbivores that feed on grains, leaves, and fruits, possess an enlarged caecum (29), retain food for extremely long time frames (12-14 days) (29), and rely on hindgut fermentation (30, 31). While the extant Family *Testudinidae* have evolved only 38-39 million years ago (Mya) (32, 33), the Order *Testudines* (with 13 other families encompassing side-necked turtles, softshell turtles, sea turtles, and others) is much older, evolving in the Late Triassic (237-201 Mya); and some of its extinct members (e.g. *Proganochelys*) were known to be herbivores, with a digestive process highly similar to extant tortoises (34).

Here, we challenge the prevailing mammalian-centric narrative of AGF distribution and evolutionary history by the identification, isolation, and characterization of ancient, deep-branching AGF taxa from the tortoise GIT. The discovery of these novel tortoise-associated AGF demonstrate that AGF evolution as a distinct fungal phylum predates the rise of mammalian herbivory post the K-Pg extinction event; previously regarded as the defining event driving *Neocallimastigomycota* evolution and establishment of herbivores-fungal nutritional symbiosis (27, 35). Finally, we assess trait evolution patterns in these novel taxa in comparison to other mammalian-associated AGF and demonstrate that massive horizontal gene transfer (HGT) events driving CAZyme expansion in the *Neocallimastigomycota* have occurred mostly in mammalian-, but not tortoise-associated AGF lineages.

## Results

### Anaerobic gut fungal diversity and community structure in tortoise

Culture-independent analysis identified the occurrence of AGF in all tortoise samples examined (n=11, Table S1). A distinct community composition pattern was observed, with sequences affiliated with three AGF genera (NY54, NY56, and NY36) being highly prevalent, representing (either individually or collectively) >90% of sequences encountered in 9/11 samples (Dataset 2, Figure 1a). Candidate genus NY54 was the most ubiquitous being identified in all tortoise samples, as well as the most abundant making up the >90% of the AGF community in 6 samples (Pancake, Impressed, Egyptian, Indian star, one Galapagos, and one Sulcata tortoises) and >50% of the AGF community in one sample (Burmese star tortoise). Candidate genus NY56 was encountered only in 3/11 samples, and in only one of these (the Texas tortoise) it constituted >90% of the AGF community, while making up only a minor fraction (<1%) in the other two samples. Similarly, candidate genus NY36 was less ubiquitous, being only encountered in two samples, and constituting >90% of the community in one sample). In contrast to their collective abundance in tortoise fecal samples, two out of the three tortoise-associated genera (NY36 and NY56) were seldom identified in reference mammalian fecal samples, while the third genus (NY54) exhibited a higher level of occurrence (Figure 1b). Regardless of their observed pattern of ubiquity, the three tortoise-associated genera constituted an extremely minor component of the community in mammalian samples, when identified (Figure 1b).

**Figure 1.**
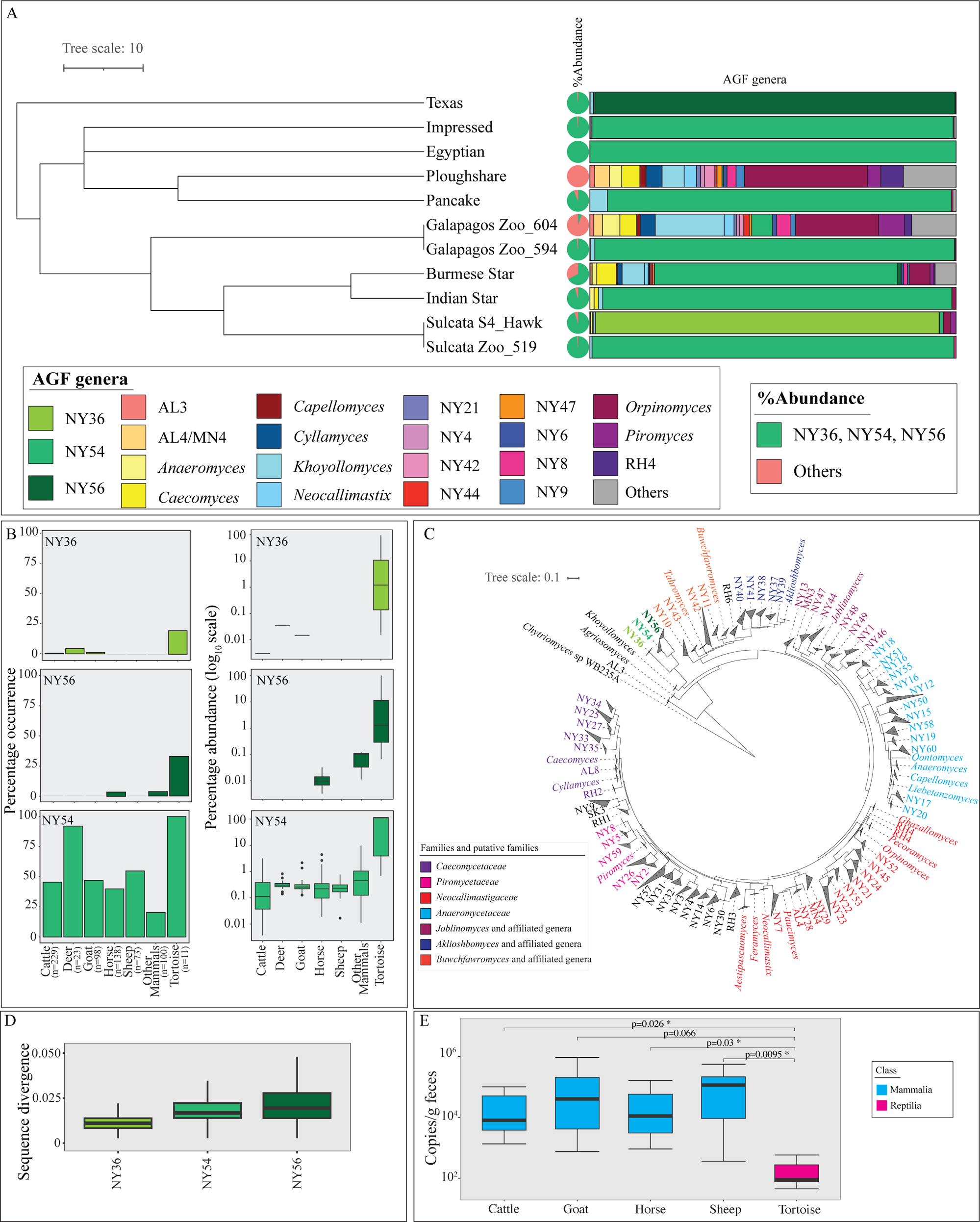
AGF diversity and community structure in Tortoises (A) Community compoisition in 11 tortoise fecal samples studied. The Tortoise phylogenetic tree downloaded from timetree.org The pie chart to the right shows the total percentage abundance of the three tortoise-affiliated genera (NY54, NY36, and NY56) (green) versus other AGF genera (peach). AGF community composition for each tortoise sample is shown to the right as colored columns corresponding to the legend key. (B) Percentage occurrence (left) and percentage abundance (right) of the three tortoise affiliated genera in previously studied cattle, white-tail deer, goat, horse, sheep, and other mammals (27), as well as in the 11 tortoise samples studied. The number of individuals belonging to each animal species is shown on the X-axis. Color code follows the key in A. (C) Maximum likelihood phylogenetic tree constructed from the alignment of the D1/D2 region of the LSU rRNA genes and highlighting the position of the three tortoise affiliated genera in relation to all previously reported cultured and uncultured AGF genera. Genera are color coded by family or putative family and the three tortoise affiliated genera are shown in green boldface. (D) Distribution of sequence divergence within each genus. (E) AGF load (determined using qPCR and expressed as copy number/g fecal sample) in the 11 tortoise samples studied here in comparison to ten individual cattle, goats, sheep, and horses selected. Significance is shown above the boxplots and correspond to Student t-test p-value).

Phylogenetic analysis using the D2 domain of the LSU rRNA placed NY54, NY56, and NY36 as three distinct, deeply-branching lineages within the *Neocallimastigomycota* tree (Figure 1c), with the closest relative being the genus *Khoyollomyces* (Figure 1c), recently identified as the earliest evolving AGF genus (36). Identical sequences across various tortoises were identified for each of the three genera (Figure S1) suggesting ready cross-host colonization. The observed low intra-genus sequence divergence estimates for all three genera suggests an extremely low level of speciation (Figure 1d). Quantitative PCR was conducted to estimate the abundance of AGF in tortoise fecal samples. AGF load (expressed as ribosomal copy number per gram feces) was invariably low in all tortoise fecal samples examined. Loads were much higher in AGF canonical mammalian hosts, e.g. cattle, sheep, goat, and horse (Figure 1e).

Assessment of alpha diversity patterns indicated that tortoises harbored a significantly less diverse AGF community when compared to placental mammals (p-value <0.04) (Figure 2a). Community assessment using PCoA constructed using phylogenetic similarity-based weighted Unifrac confirmed the clear distinction between tortoise and mammalian AGF mycobiomes (Figure 2b-c). Host class (Mammalia versus Reptilia) explained 47.6% of the variance (adonis p-value = 0.001), while the host species explained 24.6% of the community variance (adonis p-value = 0.001). DPCoA ordination plots (Figure 2d) showed that the abundance of the tortoise-associated genera NY36, NY54, and NY56, and the paucity of all other AGF was responsible for the observed pattern of community structure distinction.

**Figure 2.**
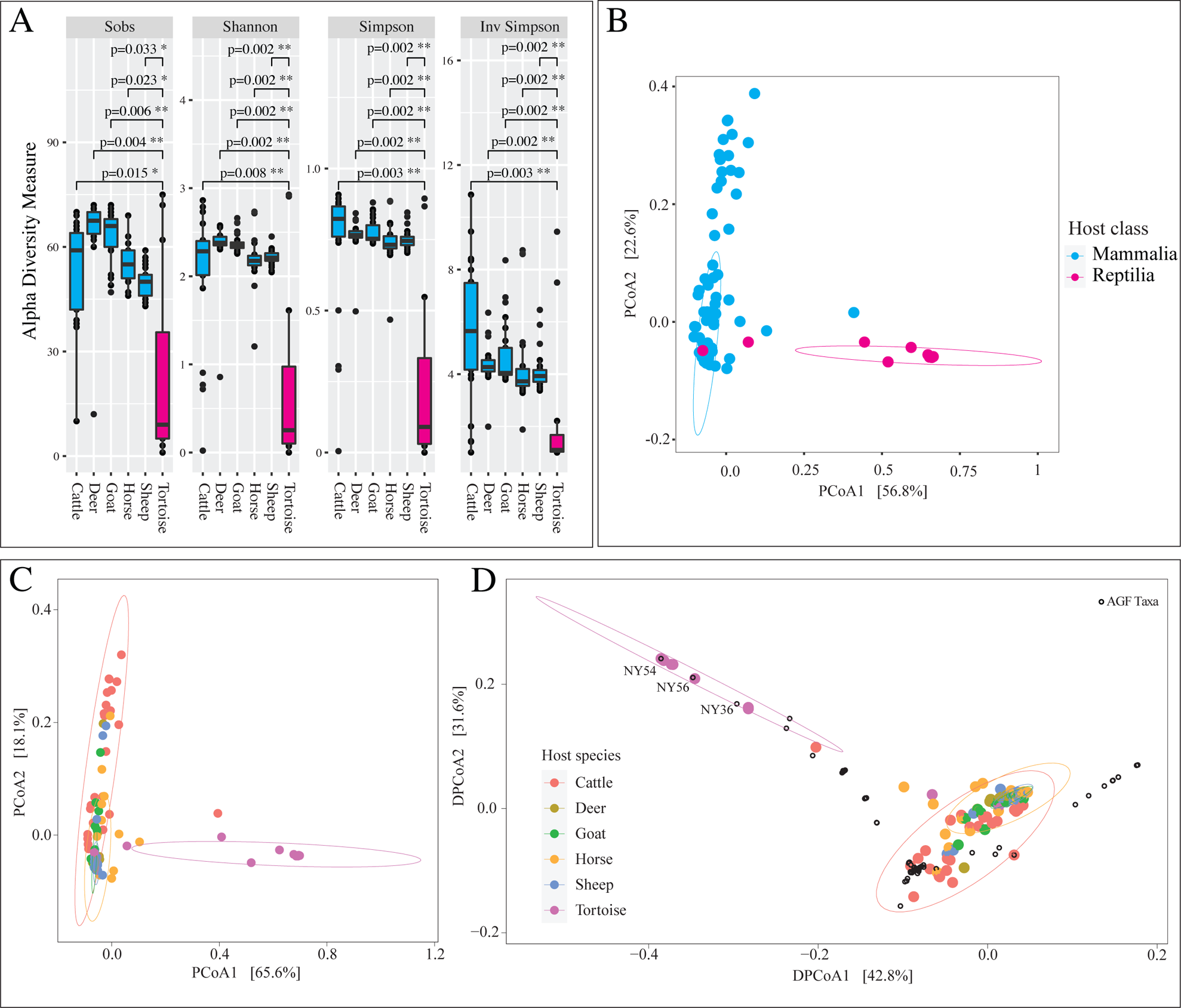
Patterns of AGF alpha and beta diversity in the 11 tortoise samples studied in comparison to a subset of mammalian hosts previously studied (27) (Dataset 1). (A) Box and whisker plots showing the distribution of 4 alpha diversity measures (observed number of genera (Sobs), Shannon, Simpson, and Inverse Simpson) for the different animal species. Results of two-tailed ANOVA for pairwise comparison of tortoise (pink) alpha diversity indices to mammals (cyan; cattle (n=25), deer (n=24), goat (n=25), horse (n=25), and sheep (n=25)). (B-C) Principal coordinate analysis (PCoA) plot based on the phylogenetic similarity-based index weighted Unifrac. The precentage variance explained by the first two axes are displayed on the axes, and ellipses encompassing 95% of variance are displayed. Samples and ellipses are color-coded by host class (B), and host species (C). Some of the circles representing tortoise samples might not be apparent due to overlap with other data points. (D) Double principal coordinate analysis (DPCoA) biplot based on the phylogenetic similarity-based index weighted Unifrac. The percentage variance explained by the first two axes are displayed on the axes, and ellipses encompassing 95% of variance are displayed. Samples and ellipses are color-coded by host species. AGF genera are shown as black empty circles and the three tortoise affiliated genera are labeled.

### Isolation of tortoise-associated AGF genera

Isolation efforts from tortoise fecal samples yielded twenty-nine different isolates (Table S2, Figure S2). Amplification and sequencing of the D1/D2 region of the LSU rRNA confirmed that the obtained isolates are identical to sequences encountered in culture-independent surveys. Representative isolates belonging to the candidate genera NY54 and NY36 have been successfully maintained and characterized as *Testudinimyces* gen. nov, and *Astrotestudinimyes* gen. nov, respectively (37). On the other hand, despite repeated successful isolation rounds, isolates belonging to candidate genus NY56 have been extremely hard to maintain as viable cultures for subsequent analysis. The names *Testudinimyces* and *Astrotestudinimyes* will henceforth be used to describe cultured strains belonging to NY54 and NY36 the manuscript.

### Phylogenomic and molecular clock timing analysis

Transcriptomics-enabled phylogenomic analysis placed isolates belonging to the genera *Testudinimyces* and *Astrotestudinimyces* as two distinct, early-branching basal lineages in the *Neocallimastigomycota* tree (Figure 3). Molecular clock timing suggests that tortoise-associated AGF (T-AGF) have evolved in the early Cretaceous, with a divergence time estimate of 112.19 Mya, with the 95% Highest Probability Density (HPD) interval at 95.94-129.98 Mya, and 104.43 Mya (95% HPD: 89.37-120.82 Mya) for *Astrotestudinimyces*, and *Testudinimyces*, respectively. Such estimates push the *Neocallimastigomycota* evolution by approximately 45 Mya, since prior efforts timed the phylum evolution at 67 Mya in the early Paleogene (27, 35), and indicate that *Neocallimastigomycota* evolution predates the K-Pg extinction event and subsequent evolution of mammalian herbivorous families (e.g. *Equidae, Bovidae, Cervidae)* as well as grasses (*Poaceae*), previously regarded as the defining process forging AGF evolution into a distinct fungal phylum (27, 35).

**Figure 3.**
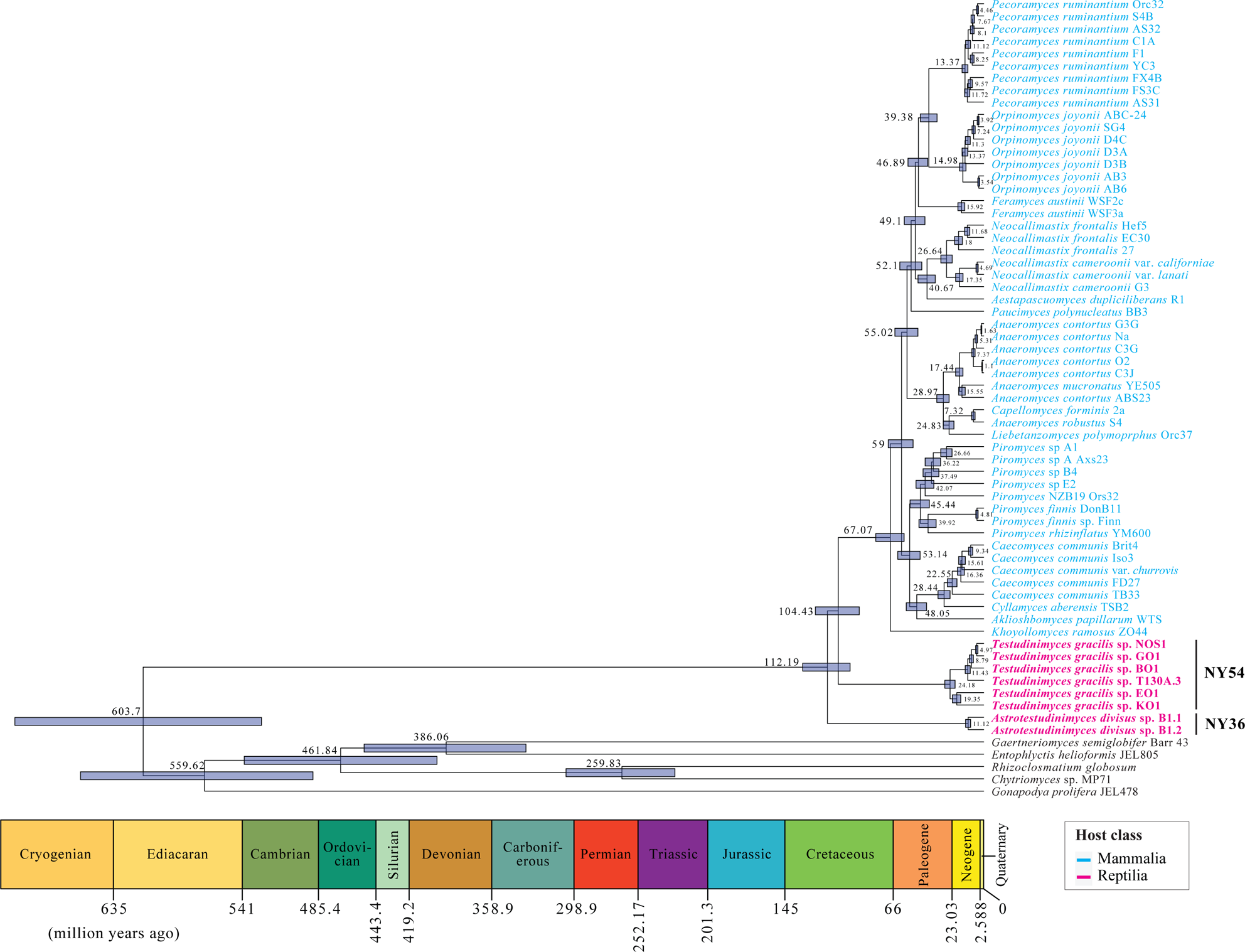
Bayesian phylogenomic maximum clade credibility (MCC) tree of *Neocallimastigomycota* with estimated divergence time. The isolate names are color coded by host class as shown in the legend. Strains belonging to the two T-AGF genera are shown in boldface and the taxa label is shown to the right. All clades above the rank of the genus are fully supported by Bayesian posterior probabilities. The 95% highest-probability density (HPD) ranges (blue bars) are denoted on the nodes, and the average divergence times are shown. Geological timescale is shown below.

### A curtailed carbohydrate active enzyme machinery in tortoise-associated AGF

Preliminary comparative transcriptomic analysis (Supp. text, Figure S3) indicated that T-AGF lack many gene clusters (n=1699) encountered in all currently available, mammalian-associated AGF (henceforth M-AGF) isolates (Figure S4). Interestingly, a significant proportion (55.13%) of these gene clusters encoded metabolic functions, with a high proportion of carbohydrate metabolism (49.63% of metabolic functions gene clusters) and, more specifically, an enrichment of Carbohydrate Active enZymes (CAZymes) (46% of carbohydrate metabolism clusters) (Figure S4). Further, representatives of the genera *Testudinimyces* (strain T130A.3) and *Astrotestudinimyces* (strain B1.1) demonstrated a slower ability to grow on (microcrystalline) cellulose, failed to grow on xylan, and exhibited a relatively more limited capacity for carbohydrates metabolism compared to reference M-AGF (Figure S5 and (37)). Such pattern strongly suggests a curtailed machinery for plant biomass degradation in tortoise-associated, compared to M-AGF isolates.

Comparative analysis demonstrated that T-AGF harbor a significantly reduced CAZyome compared to M-AGF (Student t-test p-value=0.0011), with only 0.5±0.11% of the predicted peptides assigned to a GH, CE, and PL families, compared to 1.3 ±0.61% in mammalian isolates transcriptomes (Dataset 3). Specifically, T-AGF transcriptomes harbored significantly lower number of distinct transcripts assigned to the families primarily associated with cellulose and hemicellulose metabolism, e.g. cellulase GH families GH5, GH9, the xylanase families GH10, GH11, GH16, GH45, the cellobiohydrolase families GH6, GH48, the β-glucosidase family GH3, the β-xylosidase family GH43, the α-amylase family GH13, the acetyl xylan esterases families CE1, CE2, CE4, and CE6 (Wilcoxon adjusted p-value < 0.03) (Figure 4).

**Figure 4.**
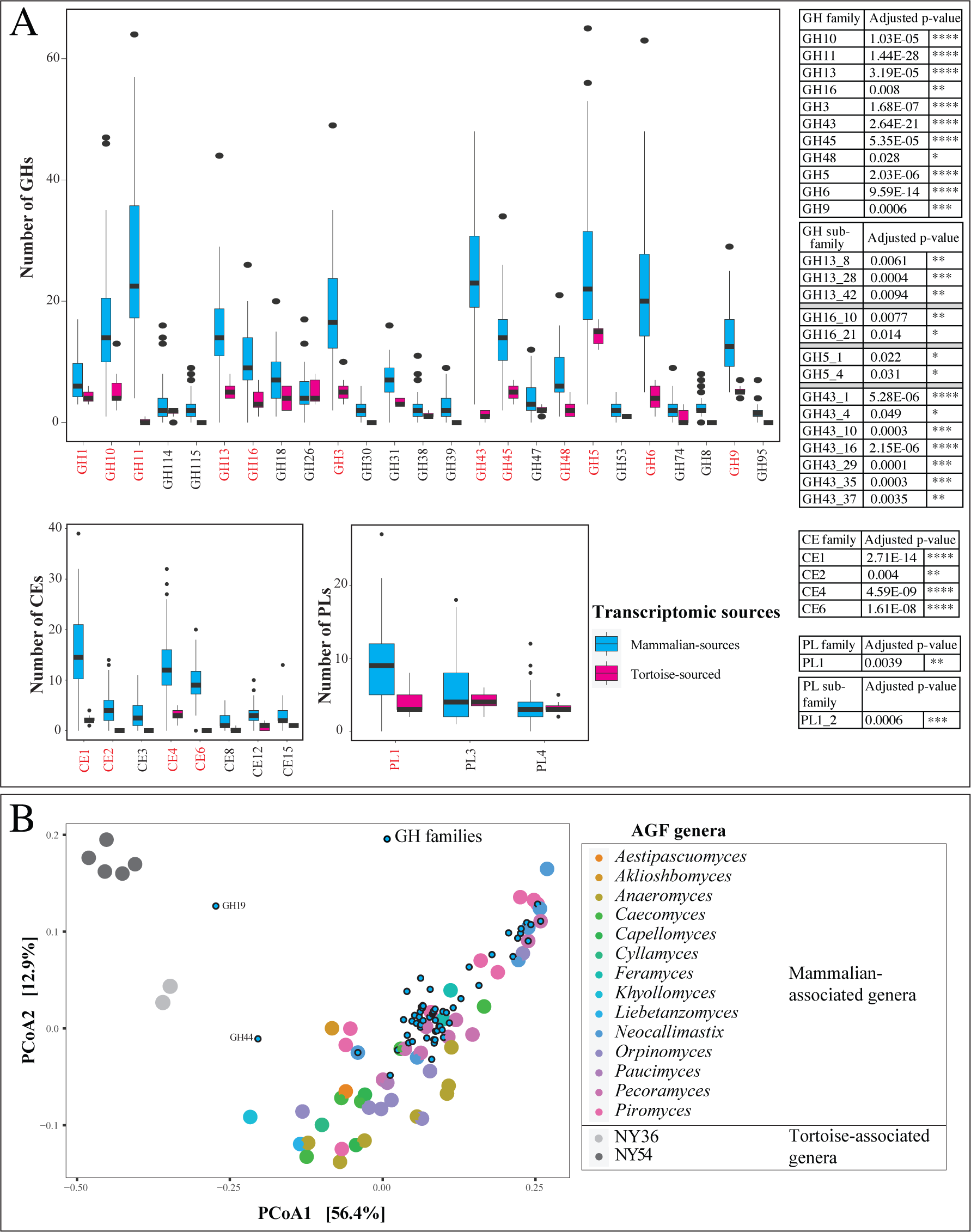
CAZyome composition difference between tortoise sourced (n=7) and mammalian sourced (n=54) strains. (A) Box and whisker plots for the distribution of the total number of GHs (top), CEs (bottom left), and PLs (bottom right) identified in the transcriptomes (mammalian sourced, cyan; tortoise sourced, pink). Only CAZy families with >100 total hits in the entire dataset are shown, and CAZy families that were significantly more abundant in mammalian versus tortoise transcriptomes are shown in red text. Wilcoxon test adjusted p-values for the significance of difference in CAZyome composition for the families in red text are shown to the right, along with the values for GH5, GH13, GH16, GH43, and PL1 sub-families. (B) Principal coordinate analysis (PCoA) biplot based on the GH families composition in the studied transcriptomes. The % variance explained by the first two axes are displayed on the axes and strains are color coded by AGF genus as shown in the figure legend to the right, while GH families are shown as smaller cyan spheres with black borders.

### Limited horizontal gene transfer in tortoise-associated AGF

Interestingly, many of the CAZyme families lacking or severely curtailed in T-AGF have previously been shown to be acquired by AGF via horizontal gene transfer (HGT) (Figure S4, (38)). To determine whether this reflects a broader pattern of sparse HGT occurrence in the entirety of T-AGF genomes, we quantified HGT occurrence and frequency in T-AGF transcriptomes. Our analysis (Table 1) identified a total of only 35 distinct HGT events (with an average of 0.16±0.05% of transcripts in the 7 sequenced T-AGF transcriptomes). This value is markedly lower than the 277 distinct HGT events previously reported from M-AGF transcriptomes (38). Interestingly, within the limited number of HGT events identified in T-AGF, the majority (30/35) were also identified in M-AGF (38); and virtually all of which (29/30) had the same HGT donor (Table 1). Such pattern indicates that most HGT events in T-AGF occurred prior to the phylum’s *Neocallimastigomycota* diversification into tortoise- and mammalian-associated lineages.

**Table 1.** HGT events identified in the two tortoise genera (B: *Astrotestudinimyces*; T: *Testudinimyces*), the affiliation of HGT donor, and distribution of the event in other AGF.

| COG/ KEGG Classification | Function imparted/ Pathway | Affiliation of donor |  | In both tortoise genera | HGT In other AGF |
| --- | --- | --- | --- | --- | --- |
|  |  | Phylum/ Class | Kingdom/ Clade |  |  |
| Cellular Processes and Signaling |  |  |  |  |  |
| [O] Posttranslational modification, protein turnover, chaperones | DsbA | Cnidaria | Metazoa | Yes | Yes |
| Information Storage and Processing |  |  |  |  |  |
| [L] Replication, recombination and repair | Methylated DNA protein-cysteine methyl transferase | Firmicutes | Bacteria | Yes | Yes |
| [J] Translation, ribosomal structure and biogenesis | GTP-binding | Mixed phyla | Bacteria | Yes | Yes |
| Metabolism |  |  |  |  |  |
| [E] Amino acid transport and metabolism | ADP-ribosyl arginine hydrolases | Firmicutes | Bacteria | Only in T | Yes |
|  | Aspartate-ammonia ligase | Bacteroidetes | Bacteria | Yes | Yes |
|  | Cysteine synthase | Firmicutes | Bacteria | Yes | Yes |
|  | Tryptophan synthase (trpB) | Verrucomicrobia | Bacteria | Only in T | Yes |
| [H] Coenzyme transport and metabolism | Dephospho CoA kinase |  | Amoebozoa | Only in T | Yes |
|  | NadB (L-aspartate oxidase) | Myxococcota | Bacteria | Yes | Yes |
| [C] Energy production and conversion | Lactate dehydrogenase and 2-hydroxyacid dehydrogenase | Firmicutes | Bacteria | Yes | Yes |
|  | Lipoamide dehydrogenase | Mixed phyla | Bacteria | Yes | Yes |
|  | Bifunctional aldehyde/alcoholDH family of Fe-ADH | Cyanobacteria | Bacteria | Yes | Yes |
| [F] Nucleotide transport and metabolism | Guanine deaminase | Firmicutes | Bacteria | Only in B | No |
|  | Thymidine kinase | Alpha-Proteobacteria | Bacteria | Only in T | Yes |
|  | Anaerobic ribonucleoside triphosphate reductase | TM6 | Bacteria | Yes | Yes |
| [Q] Secondary metabolites biosynthesis, transport and catabolism | Dehydrogenases | Firmicutes | Bacteria | Yes | Yes |
|  | Dehydrogenases | Firmicutes | Bacteria | Only in T | Yes |
| [KO] Metabolism of terpenoids and polyketides | Squalene-hopene cyclase | Ciliophora | Alveolata | Yes | Yes |
| [G] Carbohydrate transport and metabolism |  |  |  |  |  |
| Glycoside Hydrolases | GH13 | Firmicutes | Bacteria | Yes | Yes |
|  | GH1 | Firmicutes | Bacteria | Yes | Yes |
|  | GH5 | Firmicutes | Bacteria | Yes | Yes |
|  | GH43 | Firmicutes | Bacteria | Yes | Yes |
|  | GH10 | Firmicutes | Bacteria | Yes | Yes |
|  | GH11 | Fibrobacteres | Bacteria | Only in B | Yes |
|  | GH88 | Mollicutes/Firmicutes | Bacteria | Yes | Yes |
|  | GH3 | Firmicutes | Bacteria | Only in B | Yes |
|  | GH53 | Firmicutes | Bacteria | Yes | Yes |
|  | GH48 | Firmicutes and Actinobacteria | Bacteria | Only in T | Yes |
| Carbohydrate Esterases | CE1 | Firmicutes and Fibrobacteres | Bacteria | Only in B | Yes |
|  | CE15 | Bacteroidetes | Bacteria | Yes | Yes |
| Polysaccharide Lyases | PL4 | Bacteroidetes | Bacteria | Yes | Yes |
| **Poorly characterized** |  |  |  |  |  |
| *[KO] Not Included in Pathway or Brite* | Aminopeptidase (Peptidase MEROPS Family M18) | Firmicutes | Bacteria | Yes | No |
| *No COG/KEGG* | TerD_like | Firmicutes | Bacteria | Yes | No |
|  | Uncharacterized protein |  | Metazoa | Yes | No |
|  | Uncharacterized protein | Firmicutes | Bacteria | Yes | No |

Prior work has suggested the prevalence of metabolic functions in genes acquired by HGT in M-AGF (38). Such pattern held true for T-AGF, with (29/35) of the identified HGT events encoding a metabolic function. Several HGT-acquired metabolic genes in T-AGF were involved in processes enabling anaerobiosis. Specifically these genes mediated functions such as recycling reduced electron carriers via fermentation (aldehyde/alcohol dehydrogenases and d-lactate dehydrogenase for ethanol and lactate production from pyruvate), *de novo* synthesis of NAD via the bacterial pathway (L-aspartate oxidase NadB), the acquisition of the oxygen-sensitive ribonucleoside-triphosphate reductase class III (anaerobic ribonucleoside triphosphate reductase nrdD) and of squalene-hopene cyclase, catalyzing the cyclization of squalene into hopene during biosynthesis of tetrahymanol (that replaced the molecular O_2_-requiring ergosterol in the cell membranes of AGF) (Table 1). Few additional HGT-acquired metabolic genes encoded CAZymes (Table 1). However, the number of HGT-acquired CAZyme genes in T-AGF was extremely minor (13 events representing an average of 10.81±4.17% of the total CAZYome in the 7 sequenced transcriptomes) compared to the massive acquisition of CAZymes by HGT previously reported in M-AGF (a total of 72 events representing 24.62-40.41% of the overall CAZyome) (38).

### Cellulosomal production capacity in tortoise versus mammalian AGF

Anaerobic fungi produce cellulosomes, extracellular structures that function as multienzyme complexes that synergistically break down plant biomass into fermentable sugars. Many AGF-produced CAZymes localize to the cellulosomes. A non-catalytic dockerin domain (NCDD) similar to carbohydrate-binding module family 10 (CBM10) is usually associated with cellulosome-bound genes in anaerobic fungi and typically docks the enzymes to cohesin domains housed in a large scaffolding protein (scaffoldin), that in-turn anchors the entire structure to the cell wall (26). We hypothesized that the observed differences in gene content (Figure S4), CAZyme repertoire (Figure 4, Dataset 3), secretome content (Supp. text, Figure S6), and HGT frequency (Table 1) between T-AGF and M-AGF would result in a differential cellulosomal production capacity (assessed as all peptides predicted to be extracellular and harbor an NCDD, as previously suggested (39–41)). Within the transcriptomes of representatives of T-AGF genera *Testudinimyces* (strain T130A) and *Astrotestudinimyces* (strain B1.1), predicted peptides with high sequence similarity (>27.34% aa identity), and close phylogenetic affiliation (Figure 5a) to *Neocallimastigomycota* scaffoldin protein ScaA were identified (5 copies in T130A, and 38 copies in B1.1 transcriptomes equivalent to 0.03, and 0.14% of total transcripts). As well, a total of 91, and 183 transcriptome predicted peptides possessing a non-catalytic dockerin domain (NCDD) and predicted to be extracellular were identified in T130A, and B1.1, respectively (equivalent to 1.16, and 0.34% of total transcripts). NCDD-harboring predicted peptides encoded CAZymes (n=43, and 72, respectively), spore coat protein CotH (n=6, and 9, respectively), carbohydrate binding modules (n=34, and 87, respectively), expansins (n=1, and 3, respectively), and other functions including hydrolases, and phosphatases (Figure 5b). For comparative purposes, we sequenced and analyzed the transcriptome of a reference M-AGF isolate (*Orpinomyces joyonii* strain AB3). Similar to previously reported M-AGF, e.g. *Pecoramyces* (39), *Caecomyces* (42), *Piromyces*, *Neocallimastix*, and *Anaeromyces* (41), strain AB3 harbored a larger number of scaffoldin predicted peptides (n=24, equivalent to 0.146% of total transcripts), and extracellular predicted peptides possessing a NCDD (n=316, equivalent to 1.92% of total transcripts). Extracellular NCDD-harboring predicted peptides in strain AB3 encoded CAZymes (n=134), spore coat protein CotH (n=30), carbohydrate binding modules (n=118), and expansins (n=3) (Figure 5b). Further, in addition to the overall lower number of NCDD-harboring peptides in T-AGF compared to M-AGF, clear differences were also observed in the relative composition of the CAZyme component of their predicted cellulosomes. In general, an extremely minor representation of CEs and GH10 component in T-AGF cellulosomes, when compared with the M-AGF representative strain AB3, was observed (Figure 5B). On the other hand, an exclusive representation of GH45 in strain B1.1, and high and exclusive representation of PL1 in strain T130A, in comparison to strain AB3 cellulosome was observed (Figure 5c).

**Figure 5.**
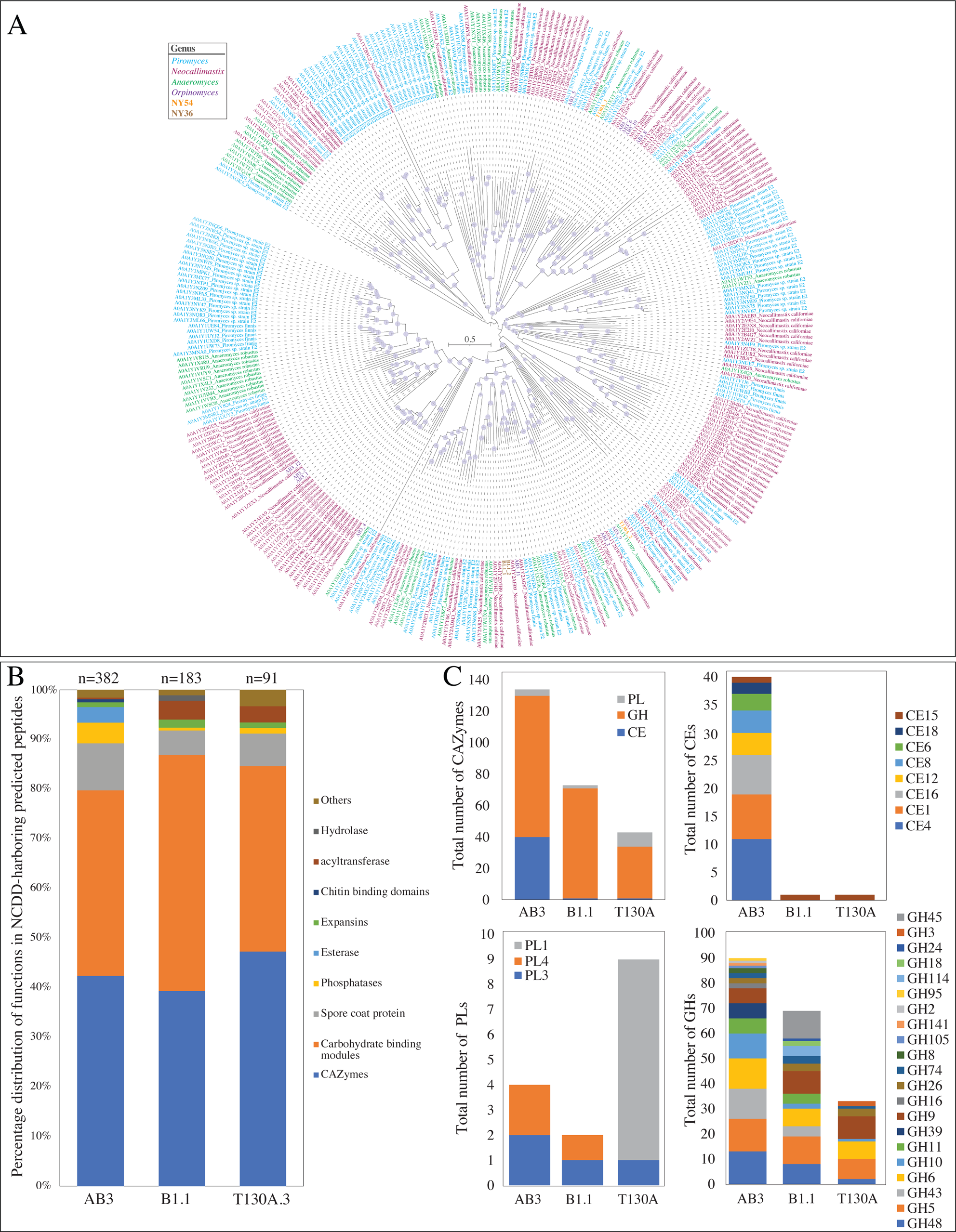
Comparative cellulosomal analysis between representatives of the two tortoise affiliated genera (genus *Astrotestudinimyces*, strain B1.1; and genus *Testudinimyces*, strain T130A) and one mammalian affiliated strain (*Orpinomyces joyonii* strain AB3). (A) Maximum likelihood mid-point rooted phylogenetic tree showing the relationship between scaffoldin ScaA protein homologues identified in *Orpinomyces joyonii* strain AB3 (12 copies denoted AB3_1 through AB3_12 and shown in purple text), *Astrotestudinimyces* strain B1.1 (2 copies denoted B1.1_and B1.1_2 and shown in brown boldface text), and *Testudinimyces* strain T130A (2 copies denoted T130A_and T130A_2 and shown in orange boldface text) in comparison to a reference set of 319 Neocallimastigomycota ScaA homologues retrieved from Uniprot. All reference ScaA homologues are shown with their Uniprot ID followed by the AGF strain name color coded by genus as shown in the legend. (B) Comparison of the percentage distribution of functions (as predicted by NCBI Conserved Domain database) encoded by cellulosomal peptides (all predicted peptides harboring a non-catalytic dockerin domain in the two tortoise affiliated genera (genus *Astrotestudinimyces*, strain B1.1; and genus *Testudinimyces*, strain T130A) and the mammalian affiliated strain (*Orpinomyces joyonii* strain AB3) and destined to the extracellular milieu (as predicted by DeepLoc)). Total numbers of peptides are shown above each column. (C) CAZyome composition of the predicted cellulosome in the three strains compared.

To confirm the translation, secretion, and cellulose-binding affinity of predicted cellulosomal proteins (scaffoldins and NCDD-containing peptides), shotgun proteomics was conducted on the total, and cellulose-bound fractions of representatives of *Astrotestudinimyces* (strain B1.1), and *Testudinimyces* (strain T130A) (Dataset 4). Of the 221, and 96 proteins predicted to be cellulosomal-bound in transcriptomics analysis of strains B1.1, and T130A, respectively, 172, and 57 proteins were identified in the proteomics dataset, confirming their translation (Dataset 5, Figure S7). Of these, 169, and 50 proteins were identified in the cellulose-bound fraction. Further, all or the majority of these proteins were identified in higher intensity in the cellulose-bound fraction (169, and 46 proteins, respectively), with intensity ratios (intensity in cellulose-bound fraction: intensity in the biomass fraction) exceeding 5 in 94%, and 90% of the proteins (Dataset 5, Figure S7).

## Discussion

Assessment of the AGF community in tortoises demonstrated that three genera (*Testudinimyces*, *Astrotestudinimyces*, and NY56) represent the majority of AGF in most samples examined (Figure 1a). The collective predominance and ubiquity of these genera is in stark contrast to their rarity and low relative abundance, when encountered, in mammalian hosts (Figure 1b, (27)). Phylogenomic and molecular clock timing analysis estimated an evolutionary time of 112 Mya, and 104 Mya for the genera *Astrotestudinimyces*, and *Testudinimyces*, respectively. Such estimates predate the evolution of all current mammalian families known to harbor AGF (43–49). More importantly, it coincides with an increased diversification in fossil records of extinct turtle lineages during the lower Cretaceous (145-100 Mya), a process spurred by an increase in climatically suitable geographic areas (50). We hence posit that tortoise-associated AGF lineages evolved in now-extinct Testudines ancestor(s) during the Middle Cretaceous, and were successfully retained throughout subsequent evolutionary events leading to the evolution of their current host (herbivorous land-dwelling tortoises, family *Testudinidae*) during the Eocene (38-39 Mya) (32, 33).

T-AGF genera were identified in 8 genera and 9 tortoise species that collectively encompass multiple feeding strategies and geographical ranges (Table S1). The observed strong pattern of tortoise-fungal phylosymbiosis indicates that T-AGF possess distinct properties enabling successful colonization and propagation in the tortoise GIT. The lower temperature optima (for *Testudinimyces*) and wider temperature range (for both *Testudinimyces* and *Astrotestudinimyces*) compared to M-AGF aids in their survival and growth in the poikilothermic (cold-blooded) tortoises, where lower and wider variation in internal temperature prevail (37). As well, the slower growth of representatives of the genus *Testudinimyces* mirrors the slower basal metabolic rate and the extremely long food retention time in tortoises (12-14 days) (29), allowing ample time for substrate colonization.

However, while a rationale for their successful establishment in the tortoise GIT over their more robust mammalian AGF counterparts could be discerned; the exact ecological role and services rendered by T-AGF to their hosts, if any, are currently unclear. Given their relatively low numbers (Figure 1e) in the ecosystem, as well as their relatively curtailed CAZyme repertoire (Figure 4 and Dataset 3), their relative contribution to substrate depolymerization in their hosts appears minor. As well, a role in oligomer conversion to monomers, followed by monomer fermentation to absorbable VFA could be postulated. Alternatively, T-AGF could be rendering ecological services unrelated to food digestion in their host, e.g., modulating community and preventing pathogenic microbes’ establishment via secondary metabolites production and niche competition, akin to the gut and skin microbiome role in colonization resistance in human (51, 52). Finally, the possibility that T-AGF are dispensable commensals, rather than indispensable symbionts could not be discounted, given their extremely low loads in the tortoise GIT (Figure 1e).

Regardless of their adaptive strategies and putative role in the tortoise GIT, our findings have important implications on our understanding of the evolutionary history of *Neocallimastigomycota*. Prior efforts based on available M-AGF taxa estimated an AGF divergence time of 67 Mya (27, 35). Such estimate post-dates the K-Pg extinction event and coincides with the evolution of mammalian AGF host families (43–49) and the associated evolutionary innovations in hosts’ alimentary tract architecture, as well as the evolution of grasses in the family *Poaceae* (53). Our results describe two distinct, deep branching lineages that independently evolved 37-45 Mya prior to these events in a non-mammalian host. The described genera hence represent the earliest known form of host-anaerobic fungal associations known to date; and demonstrate that AGF evolution predates the events previously recognized as the driving force behind forging *Neocallimastigomycota* evolution as a distinct fungal phylum. It is currently unclear whether T-AGF are the direct ancestors providing the seed for M-AGF, or whether M-AGF evolved from other yet-undiscovered extinct or extant ancestors. Nevertheless, it is important to note that despite such discovery, our results do not challenge the key role played by the rise of mammalian herbivory post the K-Pg extinction event and the evolution of mammalian families with dedicated fermentation chambers in AGF evolution. Establishment of AGF in the mammalian herbivorous gut has spurred an impressive wave of AGF family- and genus-level diversification (27), and acquisition of genes enabling efficient cellulose and hemicellulose degradation via HGT (38) to fully utilize the newly evolved grasses (family *Poaceae*) as a primary food source (35, 38). These innovations, in turn, enabled the establishment of AGF as indispensable members of the GIT tract of herbivorous mammals (23, 54–56). Indeed, most of the AGF identified diversity and biomass on earth currently resides in mammalian, rather than non-mammalian, herbivores.

Finally, comparative analysis between M- and T-AGF clearly indicates a significantly lower frequency of gene acquisition via HGT in T-AGF compared to M-AGF (Table 1). Our analysis suggests that a primary purpose of HGT in T-AGF is to enable their transition from an aerobic ancestor to an anaerobic lineage, a prerequisite for their establishment in the tortoise AGF tract. Only few (13 out of 35) HGT events were associated with improving plant degradation capacity in T-AGF lineages, which is, in turn, reflected in the slower cellulose-degradation ability and the lack of xylan degradation abilities in T-AGF taxa (Figure S5). Further, most of the HGT events identified in T-AGF in this study were also observed in M-AGF indicating ancient acquisition events prior to T-AGF and M-AGF split (Figure 3). However, these relatively few ancient HGT events were followed by a more extensive second wave of HGT-mediated gene acquisition that occurred solely in M-AGF, and was mostly responsibly for equipping M-AGF with a powerful plant biomass degradation machinery enabling their propagation, establishment, and competition in the highly competitive, bacteria and archaea dominated rumen and hindgut in mammalian herbivores (35, 38).

What prevented T-AGF genera from undergoing a similar massive acquisition of CAZymes to improve their competitive advantage in the tortoise GI tract and beyond? We provide two possible explanations for the observed deficiency. First, such difference could be niche related. Mammalian rumen and hindguts are characterized by higher temperatures, larger food intake, rapid digestion and substrate turnover, higher microbiome density and diversity and higher overall metabolic activity. Such conditions provide for a more active milieu of cells, extracellular DNA, and viruses with higher opportunities for HGT through transduction, natural uptake, and transformation. Second, such differences in HGT frequency could be related to the hyphal growth pattern of AGF taxa. T-AGF genera identified appear to be polycentric, and such taxa produce lower number of zoospores, and can depend on hyphal propagation as means of reproduction. In contrast, most M-AGF genera (16/20 genera), including earliest evolving ones (e.g. *Khoyollomyces*, *Piromyces*) are monocentric, with strict dependency on sporangial development and free zoospore release followed by encystment and growth. Given the fact that fungal zoospores are naturally competent (57, 58) and represent the most appropriate and logical stage for DNA uptake by the AGF, such differential prevalence could contribute to the observed differences in HGT between both groups.

## Materials and Methods

### Sampling, PCR amplification, amplicon sequencing, and diversity analysis

Fecal samples were obtained from animals at the Oklahoma City Zoo (Oklahoma City, Oklahoma, USA), except one Sulcate (African Spurred) tortoise sample (*Centrochelys sulcate*). DNA extraction was conducted using DNeasy Plant Pro Kit (Qiagen Corp., Germantown, MD, USA) (27). PCR amplification targeting the D2 region of the LSU rRNA using AGF-specific primers AGF-LSU-EnvS For: 5’-GCGTTTRRCACCASTGTTGTT-3’, and AGF-LSU-EnvS Rev: 5’-GTCAACATCCTAAGYGTAGGTA-3’ (59). Pooled libraries were sequenced at the University of Oklahoma Clinical Genomics Facility using the MiSeq platform. Sequence assignment to AGF genera was conducted using a two-tier approach as well as Alpha and Beta diversity estimates were conducted as described in reference (27) and in the supplementary methods.

### Quantitative PCR

We quantified AGF load in tortoise samples and compared it to samples from ten cattle, ten goats, ten sheep, and ten horses using quantitative PCR (Supplementary methods). The same primer pair (AGF-LSU-EnvS and AGF-LSU-EnvS Rev) used in the amplicon-based diversity survey described above was also used for qPCR quantification.

### Isolation of AGF from Tortoises

Isolation of AGF was conducted using established enrichment and isolation procedures in our laboratory (Supplementary methods and (25, 37)). To account for the poikilothermic (ectothermic) nature of the host, and the fact that the tortoise gut community is often exposed to lower and variable temperatures, we enriched for tortoise-associated AGF at a range of temperatures (30°C, 39°C, and 42°C).

### Transcriptomic sequencing

Transcriptomic sequencing of 7 representative tortoise-associated AGF isolates was conducted as described previously (Supplementary methods and (27, 35)). BUSCO (60) was used to assess transcriptome completeness using the fungi_odb10 dataset modified to remove 155 mitochondrial protein families as previously suggested (22).

### Phylogenomic analysis and molecular dating

Phylogenomic analysis was conducted as previously described in supplementary methods and in (27, 61) using the 7 transcriptomic datasets generated in this study, as well as 52 transcriptomic datasets from 14 AGF genera previously generated by our group (27, 35, 38), and others (22, 62–64).

### Transcriptomic gene content analysis and comparative transcriptomics

Transcriptomic datasets obtained from tortoise AGF isolates (n=7) were compared to the 52 previously generated transcriptomic datasets from mammalian AGF isolates (22, 27, 35, 38, 62–64). Gene content comparison was conducted via classification of the predicted peptides against COG, KOG, GO, and KEGG classification schemes, as well as prediction of the overall CAZyme content (Supplementary methods). To identify predicted functions that are unique to tortoise-associated or mammalian-associated AGF, predicted peptides from all 59 transcriptomes were compared in an all versus all Blastp followed by MCL clustering (Supplementary methods).

### Quantifying horizontal gene transfer (HGT)

We used an HGT detection pipeline that was previously developed and extensively validated (38) to identify patterns of HGT in AGF transcriptomic datasets. The pipeline involved a combination of BLAST similarity searches against UniProt databases, comparative similarity index (HGT index, *h_U_*), and phylogenetic analyses to identify potential HGT candidates (Supplementary methods).

### Predicted secretome in transcriptomic datasets

DeepLoc 2.0 (65) was used to predict the subcellular location of all predicted peptides from the transcriptomes (Supplementary methods). The predicted secretome was searched for the presence of scaffoldin homologues via Blastp comparison against a scaffoldin database. The predicted secretome was also searched for non-catalytic dockerin domains (NCDD) via the NCBI Batch CD-search online tool and identifying the predicted peptides with hits to the CBM_10 pfam02013 (Supplementary methods).

### Proteomics sequencing and analysis

Proteomic analysis was conducted using Liquid Chromatrography Tandem Mass Spectrometry (LC-MS/MS) on two fractions: biomass, and cellulose-bound. These fractions were obtained and purified from AGF cultures as described in the supplementary methods and (40). For LC-MS/MS, peptides were injected onto a 75 µm x 50 cm nano-HPLC column packed with 1.9-micron C18 beads (Thermo PN 164942) connected to an Easy-nLC 120 nano-HPLC system configured for two-column vented trap operation. The details regarding the MS programming are provided (Table S3). RAW files from the mass spectrometer were searched against the corresponding transcriptome predicted peptides database using the MaxQuant application (v2.0.2.0) as described in (66) and supplementary methods.

## Data availability

Illumina amplicon reads generated in this study have been deposited in GenBank under BioProject accession number PRJNA997953, and BioSample accession numbers SAMN36694530-SAMN36694536. RNA-seq reads from tortoise isolates have been deposited in GenBank under BioProject accession number PRJNA997953, and BioSample accession numbers SAMN36694608-SAMN36694614.

## Authors Contributions

Conceptualization: MSE, NHY, CJP; Methodology: MSE, NHY, CJP; Formal analysis: NHY, CJP; Investigation: CJP, DKJ, EEE, ALJ, CHM; Resources: MSE, NHY; Data Curation: NHY; Writing - Original Draft: MSE, NHY; Writing - Review & Editing: MSE, NHY, CJP, EEE, CHM; Visualization: NHY, CJP; Supervision: NHY, MSE; Project administration: NHY, MSE; Funding acquisition: NHY, MSE.

## Acknowledgments

We thank Drs. Jennifer D’Agostino, and Rebecca Snyder (Oklahoma City Zoo) for collecting and providing tortoise fecal samples.

## Supplementary methods

### Samples

Fecal samples from 11 tortoises belonging to 8 genera and 9 species were obtained between November 2020 and March 2022 (Table S1). All samples originated from animals kept at the Oklahoma City Zoo (Oklahoma City, Oklahoma, USA), except one Sulcate (African Spurred) tortoise sample (*Centrochelys sulcate*), which was obtained from a local farm near Walters, OK, USA (34°28’43.3”N 98°13’33.0”W). Specimen collection from wild tortoise populations is exceedingly difficult, since many of the sampled tortoises spp. are critically endangered, e.g., ploughshare tortoise (1), and/or have a very limited geographic range (Table S1). Freshly deposited samples were placed in 15- or 50-mL conical centrifuge tubes and transferred on ice to the laboratory, where they were stored at −20°C.

### Amplicon-based diversity surveys

DNA extraction from fecal samples was conducted using DNeasy Plant Pro Kit (Qiagen Corp., Germantown, MD, USA) according to the manufacturer’s instructions and as previously described (2). PCR amplification targeting the D2 region of the LSU rRNA utilized the DreamTaq Green PCR Master Mix (ThermoFisher, Waltham, Massachusetts), and AGF-specific primers AGF-LSU-EnvS For: 5’-GCGTTTRRCACCASTGTTGTT-3’, and AGF-LSU-EnvS Rev: 5’-GTCAACATCCTAAGYGTAGGTA-3’ (3). The primers target a ∼370 bp region of the LSU rRNA gene (corresponding to the D2 domain), hence allowing for high throughput sequencing using the Illumina MiSeq platform. Primers were modified to include the Illumina overhang adaptors. PCR reactions contained 2 µl of DNA, 25 µl of the DreamTaq 2X Master Mix (Life Technologies, Carlsbad, California, USA), 2 µl of each primer (10 µM) in a 50 µl reaction mix. The PCR protocol consisted of an initial denaturation for 5 min at 95 °C followed by 40 cycles of denaturation at 95 °C for 1 min, annealing at 55 °C for 1 min and elongation at 72 °C for 1 min, and a final extension of 72 °C for 10 min. PCR products were individually cleaned to remove unannealed primers using PureLink® gel extraction kit (Life Technologies), and the clean product was used in a second PCR reaction to attach the dual indices and Illumina sequencing adapters using Nexterra XT index kit v2 (Illumina Inc., San Diego, California). These second PCR products were then cleaned using PureLink® gel extraction kit (Life Technologies, Carlsbad, California), individually quantified using Qubit® (Life Technologies, Carlsbad, California), and pooled using the Illumina library pooling calculator (https://support.illumina.com/help/pooling-calculator/pooling-calculator.htm) to prepare 4-5 nM libraries. Pooled libraries (300-350 samples) were sequenced at the University of Oklahoma Clinical Genomics Facility using the MiSeq platform.

### Sequence data analysis

Forward and reverse Illumina reads were assembled using make.contigs command in mothur (4), followed by screening to remove sequences with ambiguous bases, sequences with homopolymer stretches longer than 8 bases, and sequences that were shorter than 200 or longer than 380 bp. Sequence assignment to AGF genera was conducted using a two-tier approach as recently described (2). Briefly, sequences were first compared by Blastn to the curated D1/D2 LSU rRNA AGF database (www.anaerobicfungi.org), and were classified as their first hit taxonomy if the percentage similarity to the first hit was > 96% and the two sequences were aligned over >70% of the query sequence length. For all sequences that could not be confidently assigned to an AGF genus by Blastn, insertion into a reference LSU tree (with representatives from all cultured and uncultured AGF genera and candidate genera) was used to assess novelty. These genus-level assignments were then used to build a taxonomy file in mothur, which was subsequently used to build a shared file using the mothur commands phylotype and make.shared. The genus-level shared file was used for all downstream analyses as detailed below.

Alpha diversity estimates (observed number of genera, Shannon, Simpson, and Inverse Simpson diversity indices) were calculated using the command estimate_richness in the Phyloseq R package (5). To compare beta diversity and community structure between AGF communities in tortoises and AGF communities in canonical mammalian hosts, the AGF genus-level shared file from the 11 tortoise samples studied here was combined with the AGF genus-level shared file from a subset of mammalian hosts previously studied (2) (Dataset 1). The subset of mammalian AGF hosts included a comparable size from each of the five most-commonly sampled and numerous mammalian hosts: cattle (n=25) (*Bos taurus*), sheep (n=25) (*Ovis aries*), goat (n=25) (*Capra hircus*), white-tail deer (n=24) (*Odocoileus virginianus*), and horse (n=25) (*Equus caballus*) generated in a prior study (2) that utilized the same DNA extraction, D2 LSU region amplification, and Illumina sequencing chemistry used in this study (Dataset 1). With this combined AGF genus-level shared file, we used the ordinate command in the Phyloseq R package to calculate weighted Unifrac beta diversity indices, and used the obtained pairwise values to construct ordination plots (both PCoA and NMDS) using the function plot_ordination in the Phyloseq R package.

### Quantitative PCR

We quantified total AGF load in the 11 tortoise samples and compared it to a subset of the samples from ten cattle, ten goats, ten sheep, and ten horses (sample names in red text in Dataset 1) using quantitative PCR. The same primer pair (AGF-LSU-EnvS and AGF-LSU-EnvS Rev) used in the amplicon-based diversity survey described above was also used for qPCR quantification. The 25-μl PCR reaction volume contained 1 μl of extracted DNA, 0.3 μM of primers AGF-LSU-EnvS primer pair, and SYBR GreenER™ qPCR SuperMix for iCycler™ (ThermoFisher, Waltham, MA, USA). Reactions were run on a MyiQ thermocycler (Bio-Rad Laboratories, Hercules, CA). The reactions were heated at 95°C for 8.5 min, followed by 40 cycles, with one cycle consisting of 15 sec at 95°C and 1 min at 55°C. A pCR 4-TOPO or pCR-XL-2-TOPO plasmid (ThermoFisher, Waltham, Massachusetts) containing an insert spanning ITS1-5.8S rRNA-ITS2-D1/D2 region of 28S rRNA from a pure culture strain was used as a positive control, as well as to generate a standard curve. The efficiency of the amplification of standards (E) was calculated from the slope of the standard curve and was found to be 0.89.

### Isolation of AGF from Tortoises

Isolation of AGF from fecal samples of tortoises was conducted using established enrichment and isolation procedures in our laboratory (6, 7). A sequence-guided strategy, where samples with the highest proportion of novel, yet-uncultured AGF taxa were prioritized, was employed. To account for the poikilothermic (ectothermic) nature of the host, and the fact that the tortoise gut community is often exposed to lower and variable temperatures, we enriched for tortoise-associated AGF at a range of temperatures (30°C, 39°C, and 42°C). Finally, the rumen fluid medium used for enrichment and isolation was amended with cellobiose (RFC medium) in addition to an insoluble substrate (switchgrass) and antibiotics (50 µg ml^−1^ chloramphenicol, 20 µg ml^−1^ streptomycin, 50 µg ml^−1^ penicillin, 50 µg ml^−1^ kanamycin, and 50 µg ml^−1^ norfloxacin).

### Transcriptomic sequencing

Transcriptomic sequencing of 7 representative tortoise-associated AGF isolates (*Testudinimyces* strains BO1, EO1, GO1, NOS1, T130A.3, and *Astrotestudinimyces* strains B1.1, and B1.2) was conducted as described previously(2, 8). Briefly, biomass from cultures grown in RFC medium was vacuum filtered and used for total RNA extraction using an Epicentre MasterPure Teast RNA purification kit (Epicentre, Madison, WI) according to manufacturer’s instructions. RNA-seq was conducted on an Illumina HiSeq2500 platform using 2 × 150 bp paired-end library at the Oklahoma State University Genomics and Proteomics Core Facility. RNA-seq reads were quality trimmed and *de novo* assembled using Trinity (v2.14.0) and default parameters. Assembled transcripts were clustered using CD-HIT (9) (identity parameter of 95% (–c 0.95)) to identify unigenes. Following, peptide and coding sequence prediction was conducted on the unigenes using TransDecoder (v5.0.2) with a minimum peptide length of 100 amino acids (https://github.com/TransDecoder/TransDecoder). BUSCO (10) was used to assess transcriptome completeness using the fungi_odb10 dataset modified to remove 155 mitochondrial protein families as previously suggested (11).

### Phylogenomic analysis and molecular dating

Phylogenomic analysis was conducted as previously described (2, 12) using the 7 transcriptomic datasets generated in this study, as well as 52 transcriptomic datasets from 14 AGF genera previously generated by our group (2, 8, 13), and others (11, 14–16), in addition to 5 outgroup *Chytridiomycota* genomes (*Chytriomyces* sp. strain MP 71, *Entophlyctis helioformis* JEL805, *Gaertneriomyces semiglobifer* Barr 43, *Gonapodya prolifera* JEL478, and *Rhizoclosmatium globosum* JEL800) to provide calibration points. The final alignment file included 88 genes that were gap free and comprising more than 150 nucleotide sites. This refined alignment was further grouped into 20 partitions, each assigned with an independent substitution model, suggested by a greedy search using PartitionFinder v2.1.1. All partition files, along with their corresponding models, were imported into BEAUti v1.10.4 for conducting Bayesian and molecular dating analyses. Calibration priors were set as previously described (8) including a direct fossil record of *Chytridiomycota* from the Rhynie Chert (407 Mya) and the emergence time of *Chytridiomycota* (573 to 770 Mya as 95% HPD)). The Birth-Death incomplete sampling tree model was employed for interspecies relationship analyses. Unlinked strict clock models were used for each partition independently. Three independent runs were performed for 30 million generations each. Tracer v1.7.1 (17) was used to confirm that sufficient effective sample size (ESS>200) was obtained after the default burn-in (10%). The maximum clade credibility (MCC) tree was compiled using TreeAnnotator v1.10.4 (18).

### Transcriptomic gene content analysis and comparative transcriptomics

Transcriptomic datasets obtained from tortoise AGF isolates (n=7) were compared to the 52 previously generated transcriptomic datasets from mammalian AGF isolates (2, 8, 11, 13–16). Gene content comparison was conducted via classification of the predicted peptides against COG, KOG, GO, and KEGG classification schemes, as well as prediction of the overall CAZyme content. COG and KOG classifications were carried out via Blastp comparisons of the predicted peptides against the most updated databases downloaded from NCBI ftp server (https://ftp.ncbi.nih.gov/pub/COG/COG2020/data/ for COG 2020 database update, and https://ftp.ncbi.nih.gov/pub/COG/KOG/ for KOG database). GO annotations were obtained by first running Blastp comparisons of the predicted peptides against the SwissProt database. The first SwissProt hit of each peptide was then linked to a GO number by awk searching the file idmapping_selected.tab available from the Uniprot ftp server (https://ftp.uniprot.org/pub/databases/uniprot/current_release/knowledgebase/idmapping/idmapping_selected.tab.gz). GO numbers corresponding to the first hits were then linked to their GO aspect (one of: Molecular function, Cellular component, or Biological process) by awk searching the file “goa_uniprot_all.gaf” available from GOA ftp site (ftp://ftp.ebi.ac.uk/pub/databases/GO/goa/UNIPROT). KEGG classification was conducted by running GhostKOALA (19) search on the predicted peptides. The overall CAZyme content was predicted using run_dbcan4 (https://github.com/linnabrown/run_dbcan), the standalone tool of the dbCAN3 server (http://bcb.unl.edu/dbCAN2/) to identify GHs, PLs, CEs, AAs, and CBMs in the 7 transcriptomic datasets.

To identify predicted functions that are unique to tortoise-associated or mammalian-associated AGF, predicted peptides from all 59 transcriptomes were compared in an all versus all Blastp followed by MCL clustering. Clusters obtained were then examined to identify these clusters that are unique to both tortoise isolates, unique to one of them, or present in mammalian associated genera but absent from both tortoise genera (thereafter “GroupD” clusters). KEGG classifications of predicted peptides belonging to each of these groups were then compared.

### Quantifying horizontal gene transfer (HGT)

We implemented an HGT detection pipeline that was previously developed and extensively validated (13) to identify patterns of HGT in the 7 tortoise-associated AGF transcriptomic datasets. The pipeline involved a combination of BLAST similarity searches against UniProt databases (downloaded January 2023), comparative similarity index (HGT index, *h_U_*), and phylogenetic analyses to identify potential HGT candidates. The downloaded Uniprot databases encompassed *Bacteria*, *Archaea*, Viruses, *Viridiplantae*, *Opisthokonta-Chaonoflagellida*, *Opisthokonta-Metazoa*, the *Opisthokonta-Nucleariidae* and *Fonticula* group, all other *Opisthokonta*, and all other non-*Opisthokonta*, non-*Viridiplantae Eukaryota*. Each predicted peptide from the 7 tortoise isolates transcriptomic datasets was searched against each of these databases, as well as against the *Opisthokonta-Fungi* (without *Neocallimastigomycota* representatives). Candidates with a Blastp bit-score against a nonfungal database that was >100 and an HGT index *h_U_* that was ≥30 were further evaluated via phylogenetic analysis to confirm HGT occurrence, and to determine the potential donor. All potential candidates were first clustered using CD-HIT and 95% similarity cutoff. Representatives of each cluster were then queried against the nr database using web Blastp once against the full nr database and once against the *Fungi* (taxonomy ID 4751) excluding the *Neocallimastigomycetes* (taxonomy ID 451455) with an E value below e^−10^. The first 100 hits obtained using these two Blastp searches were downloaded and combined in one FASTA file that was then combined with the AGF representative sequences and aligned using MAFFT multiple sequence aligner, and the alignment was subsequently used to construct maximum likelihood phylogenetic trees using FastTree. At this level, candidates that showed a nested phylogenetic affiliation that was incongruent to organismal phylogeny with strong bootstrap supports were deemed horizontally transferred.

### Predicted secretome in transcriptomic datasets

To identify the predicted secretome, DeepLoc 2.0 (20) was used to predict the subcellular location of all predicted peptides from the transcriptomes of a representative of two different tortoise-associated AGF genera (strain T130A.3 and B1.1), as well as one representative of mammalian-associated AGF genera (*Orpinomyces joyonii* strain AB3). All transcripts encoding peptides predicted to be extracellular (henceforth predicted secretome) were then subjected to run_dbcan4 (https://github.com/linnabrown/run_dbcan) to identify GHs, PLs, and CEs in the predicted secretome. In addition, the predicted secretome was searched for the presence of scaffoldin homologues via Blastp comparison against a scaffoldin database (319 proteins downloaded June 2023 from Uniprot and created by searching the UniprotKB for Scaffoldin and filtering the output by taxonomy using taxid Neocallimastigomycetes [451455]). Finally, the predicted secretome was also searched for the presence of non-catalytic dockerin domains (NCDD) via the NCBI Batch CD-search online tool and identifying the predicted peptides with hits to the CBM_10 pfam02013. All predicted extracellular peptides with NCDD were further subjected to run_dbcan4 to identify co-existing GH, PL, and CE domains.

### Proteomics sequencing and analysis

In addition to secretome prediction from transcriptomic datasets, we conducted proteomic analysis on the same tortoise-associated AGF described above (*Testudinimyces gracilis* strains T130A.3 and *Astrotestudinimyces divisus* strains B1.1). Proteomic analysis was conducted on two fractions: biomass, and cellulose-bound. Briefly, cultures were grown in RFC media until mid-exponential phase (typically 3 days for *Astrotestudinimyces* strain B1.1, and 1 week for *Testudinimyces* strain T130A). Biomass fraction was first collected by centrifugation (3220 xg for 10 minutes at 4°C). The cellulosomal fraction (in the supernatant) was separated using cellulose precipitation as previously described (14);Ali, 1995 #230}. Briefly, the supernatant pH was adjusted to 7.5, followed by adding cellulose (Sigmacell type 50) (0.4% w/v) and gently stirring at 4°C for 2 hours. Low-speed centrifugation (3220 xg for 10 minutes at 4°C) was then used to separate the cellulosomal (pellet) fraction. Proteins bound to cellulose in the cellulosomal fraction (pellet) were then eluted in water by agitation at room temperature for 1 hour, followed by removal of cellulose by centrifugation. The soluble eluates were collected by centrifugation, and frozen at −80°C. For both fraction (biomass, and cellulose-bound), the frozen samples were dried by vacuum centrifugation. Dried samples were redissolved for 30 min at RT in reducing buffered guanidine (6M guanidine HCl, 0.1M Tris HCl, tris(2-carboxyethyl)phosphine, pH 8.5). Debris were removed by centrifugation, and the solutes were alkylated by adding iodoacetamide to 10 mM and incubation for 30 min in the dark at RT. The alkylation reactions were then digested with trypsin using a filter aided sample preparation (FASP) approach (21). For FASP, the samples were loaded into 30-kDa spin filter devices (Sigma®), and subjected to three buffer exchanges into 8 M urea, 0.1M TrisHCl, pH 8.5, followed by three additional buffer exchanges into digestion buffer (100 mM TrisHCl, pH 8.5). For the final buffer exchange, samples were concentrated to ∼10 µl in digestion buffer, followed by dilution with 75 µl of digestion buffer containing 0.75 µg of trypsin/LysC mix (Promega). Reactions were digested overnight at 37°C, and the trypsinolysis products were recovered by centrifugation of the FASP device. Recovered peptides were desalted using centrifugal devices loaded with C18 resin following the manufacturer’s recommendations (HMMS18R, The Nest Group). The desalted peptides were frozen and dried by vacuum centrifugation, and redissolved in 0.1% aqueous formic acid immediately prior to analysis by LC-MS/MS.

For LC-MS/MS, peptides were injected onto a 75 µm x 50 cm nano-HPLC column packed with 1.9-micron C18 beads (Thermo PN 164942) connected to an Easy-nLC 120 nano-HPLC system configured for two-column vented trap operation. Peptides were separated by gradient chromatography using 0.1% aqueous formic acid as mobile phase A and 80:20:0.1 acetonitrile/water/formic acid as mobile phase B. Peptides separations used a gradient of 4 – 32% mobile phase B delivered over a period of 120 minutes. Eluted peptides were ionized in a Nanospray Flex Ion source using stainless steel emitters (Thermo). Peptide ions were analyzed in a quadrupole-Orbitrap mass spectrometer (Fusion model, Thermo) using a “high/low” “top-speed” data-dependent MS/MS scan cycle that consisted of an MS1 scan in the Orbitrap sector, ion selection in the quadrupole sector, high energy collision in the ion routing multipole, and fragment ion analyses in the ion trap sector. The details regarding the MS programming are provided (Table Sx).

RAW files from the mass spectrometer were searched against the corresponding transcriptome predicted peptides database using the MaxQuant application (v2.0.2.0, (22)). Searches utilized MaxQuant defaults, supplemented with two additional peptide modifications: deamidation of N/Q residues, and Q cyclization to pyro-glutamate. The MaxQuant “match between runs” algorithm was not used. Sequences for reversed-sequence decoy proteins and common contaminants proteins were utilized for the database searches, but were removed from the final MaxQuant protein results.

## Supplementary results

### Comparative transcriptomic analysis of tortoise- and mammalian-affiliated AGF isolates

Comparative analysis of the 7 transcriptomes originating from tortoise AGF isolates to the 52 mammalian sourced AGF transcriptomes revealed similar distinct transcript numbers, albeit with significantly shorter average length (Figure S3a, Student t-test p-value= 0.0398), and significantly higher AT content (Figure S3a, Student t-test p-value= 0.00025). The overall GO, COG, KOG, and KEGG composition did not vary by the source of isolation (mammalian versus tortoise) (Figure S3b). However, comparative gene content analysis identified distinct transcripts that are unique to both tortoise isolates (Clusters GroupA; n=384 functional clusters), unique to one of them (i.e., present in NY36 but not NY54 or the mammalian affiliated AGF isolates transcriptomes (Clusters GroupB; n=4231 functional clusters), and vice versa (Clusters GroupC; n=3199 functional clusters), or present in mammalian affiliated AGF isolates but absent from both tortoise affiliated AGF isolates (Clusters GroupD; n=1699 functional clusters). KEGG analysis of these functional clusters revealed that 66.43-72.31% of the functions unique to both (GroupA) or either (GroupB and GroupC) of the tortoise clades were related to genetic information processing, environmental information processing and cellular processes, while only 14.09-29.93% were related to metabolism. On the other hand, clusters that were unique to the mammalian isolates (GroupD) were mainly associated with a metabolic function (53.13%) (Figure S4). Further analysis of the clusters unique to the mammalian isolates revealed that most of the metabolic functions were related to carbohydrate metabolism (49.63% of metabolic functions) (Figure S4B), which in turn were enriched in CAZymes (46.01% of carbohydrate metabolism) geared towards lignocellulose degradation (13 GH families, and 1 PL family) (Figure S4C). Interestingly, 23.35% of GroupD clusters were previously shown to be acquired by horizontal gene transfer (13). Also, the majority of the GH families enriched in the mammalian isolate transcriptomes were previously shown to be completely (red in Figure S4C), or partly (blue in Figure S4C) acquired via HGT (13). Such pattern led us to postulate that the observed curtailed capacity for substrate degradation observed in tortoise-affiliated AGF (Figure S5) is due to their possession of a limited extracellular enzyme machinery compared to mammalian-affiliated AGF, and that such limited machinery is mostly due to the lack of widespread HGT events previously observed in mammalian AGF (13).

### Limited horizontal gene transfer in tortoise-associated AGF

To quantify the contribution of HGT, or lack thereof, to shaping tortoise AGF transcriptomes, we employed the HGT detection pipeline previously utilized for mammalian AGF transcriptomes. Using this pipeline, only 35 distinct HGT events (with an average of 0.16±0.05% of transcripts in the 7 sequenced T-AGF transcriptomes) (Table 1). This value is markedly lower than the 277 events previously reported from mammalian sourced AGF transcriptomes (13). In addition to the relative paucity of HGT events, two interesting patterns emerged. First, thirty of the 35 events identified in tortoise-sourced transcriptomes as horizontally transferred were also previously reported in mammalian AGF as horizontally transferred (13), with only 5 events exclusive for the tortoise sourced transcriptomes. These shared HGT events between mammalian and tortoise AGF also share the identity of the donor, with a bacterial origin for 27/30 shared HGT events and 26 of these 27 bacterial events sharing the same donor phylum. In addition, out of the four HGT events in tortoise-sourced transcriptomes with eukaryotic origin, two shared the same donor with mammalian sourced transcriptomes. These results imply the occurrence of ancient horizontal gene transfer events that were retained post-diversification of the mammalian AGF genera. Secondly, the majority of HGT events (29/35) encoded a metabolic function contributing to survival in the anaerobic gut. These functions included recycling reduced electron carriers via fermentation (aldehyde/alcohol dehydrogenases and d-lactate dehydrogenase for ethanol and lactate production from pyruvate), *de novo* synthesis of NAD via the bacterial pathway, the acquisition of the oxygen-sensitive ribonucleoside-triphosphate reductase class III and of squalene-hopene cyclase, catalyzing the cyclization of squalene into hopene during biosynthesis of tetrahymanol (that replaced the molecular O_2_-requiring ergosterol in the cell membranes of AGF). While the tortoise AGF CAZYome was significantly curtailed (Figure 5), some of the HGT events identified in tortoise-sourced transcriptomes involved CAZyme families acquired from bacterial members of the gut (Firmicutes, Bacteroidetes, and Fibrobacteres). However, the number of HGT-acquired CAZyme genes in T-AGF was extremely minor (13 events representing an average of 10.81±4.17% of the total CAZYome in the 7 sequenced transcriptomes) compared to the massive acquisition of CAZymes by HGT previously reported in M-AGF (a total of 72 events representing 24.62-40.41% of the overall CAZyome) (13).

### Tortoise-affiliated AGF secretome

To examine whether the curtailed CAZyome in tortoise isolates is part of a broader pattern of an overall curtailed secretome, we compared the predicted secretome (transcriptome predicted peptides destined to the extracellular milieu as predicted by DeepLoc) of a mammalian AGF isolate, *Orpinomyces joyonii* strain AB3, to these of the tortoise isolates B1.1, and T130A (each representing one of the AGF affiliated genera NY36, and NY54, respectively). Results showed that a smaller percentage (6.98-7.02%) of tortoise isolates predicted peptides were extracellular (using DeepLoc), as opposed to 11.49% of the predicted peptides of the mammalian isolate (Figure S6A). The mammalian isolate predicted secretome was slightly more enriched in carbohydrate metabolism (Figure S6C). Further, only 11.98-12.69% of the predicted secretome was affiliated with a CAZyme family in the tortoise affiliated strains, as opposed to 18.28% in the mammalian sourced isolate *Orpinomyces joyonii* strain AB3 (Figure S6D), with a slightly different CAZYome composition (Figure S6E).

## Supplementary Tables

**Table S1.**
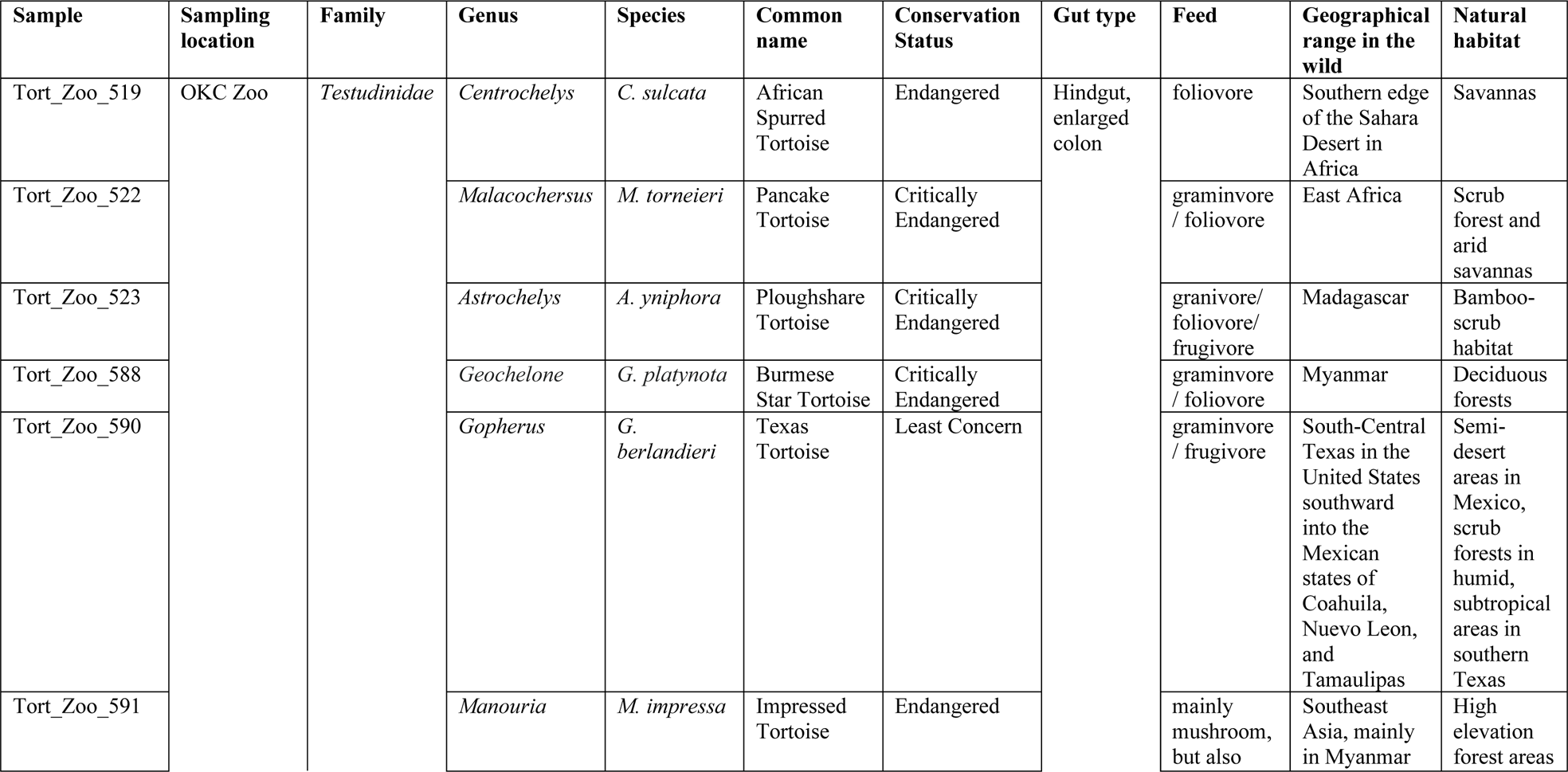
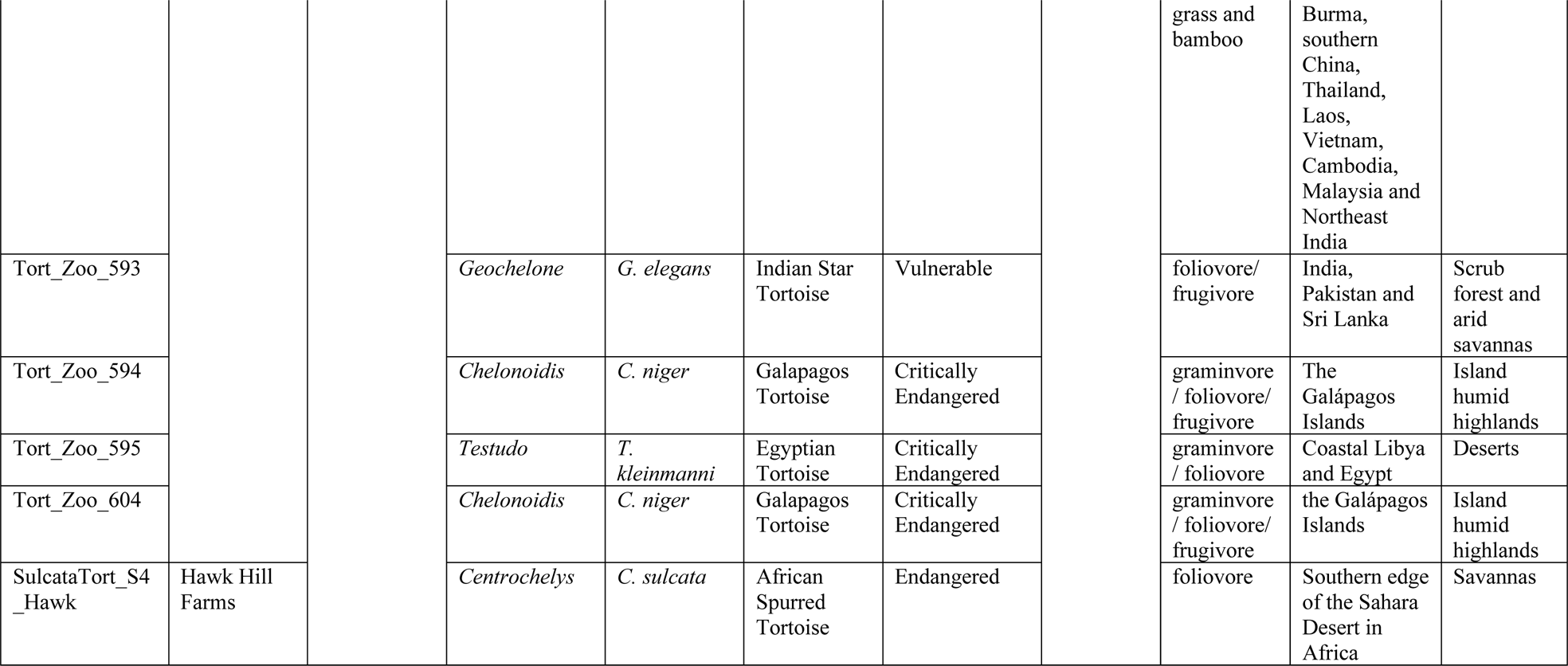
The 11 tortoise samples studied here, along with their sampling locations. All tortoises belonged to the same family but were distributed into 8 genera and 9 species. Information on conservation status was from (23), while information on feed, geographical range and natural habitat was obtained from the US Fish and Wildlife Service website (https://www.fws.gov/).

**Table S2.** List of isolates obtained from eight tortoise species, the sampling location, and the candidate genus they belong to. Isolate names in boldface have been used for transcriptomic sequencing. Isolates belonging to candidate genera NY54 and NY36 have been formally characterized and named in (7). The isolate belonging to candidate genus NY56 has been extremely hard to maintain as a viable culture for subsequent analysis.

| Tortoise species | Location | Isolate Names | Identity |
| --- | --- | --- | --- |
| Egyptian tortoise (*Testudo kleinmanni*) | Oklahoma City Zoo | **E01** | NY54 |
| Galápagos tortoise (*Chelonoidis niger*) | Oklahoma City Zoo | G01, **G01.1**, G01.2, G01.3, G01.4, G01.5 |  |
| Indian star tortoise (*Geochelone elegans*) | Oklahoma City Zoo | N0S0.1, N0S0.3, **N0S1.1**, N0S1.3, N0S2 |  |
| Pancake tortoise (*Malacochersus tornieri*) | Oklahoma City Zoo | P00, **P01** |  |
| Ploughshare tortoise (*Astrochelys yniphora*) | Oklahoma City Zoo | T030A, T030A.3, T130A, **T130A.3**, T230A, T230A.3, T0395, T0397, T0399 |  |
| Burmese star tortoise (*Geochelone platynotan*) | Oklahoma City Zoo | B01.1, B01.2, B01.3, **B1.1**, **B1.2**, B0.3 | NY36 |
| Sulcata tortoise (*Centrochelys sulcata*) | Hawk Hill Farms | S1.1, S1.2, S1.3, S1.4, S09, S19, S29 |  |
| Texas Tortoise (*Gopherus berlandieri*) | Oklahoma City Zoo | X1 | NY56 |

**Table S3:**
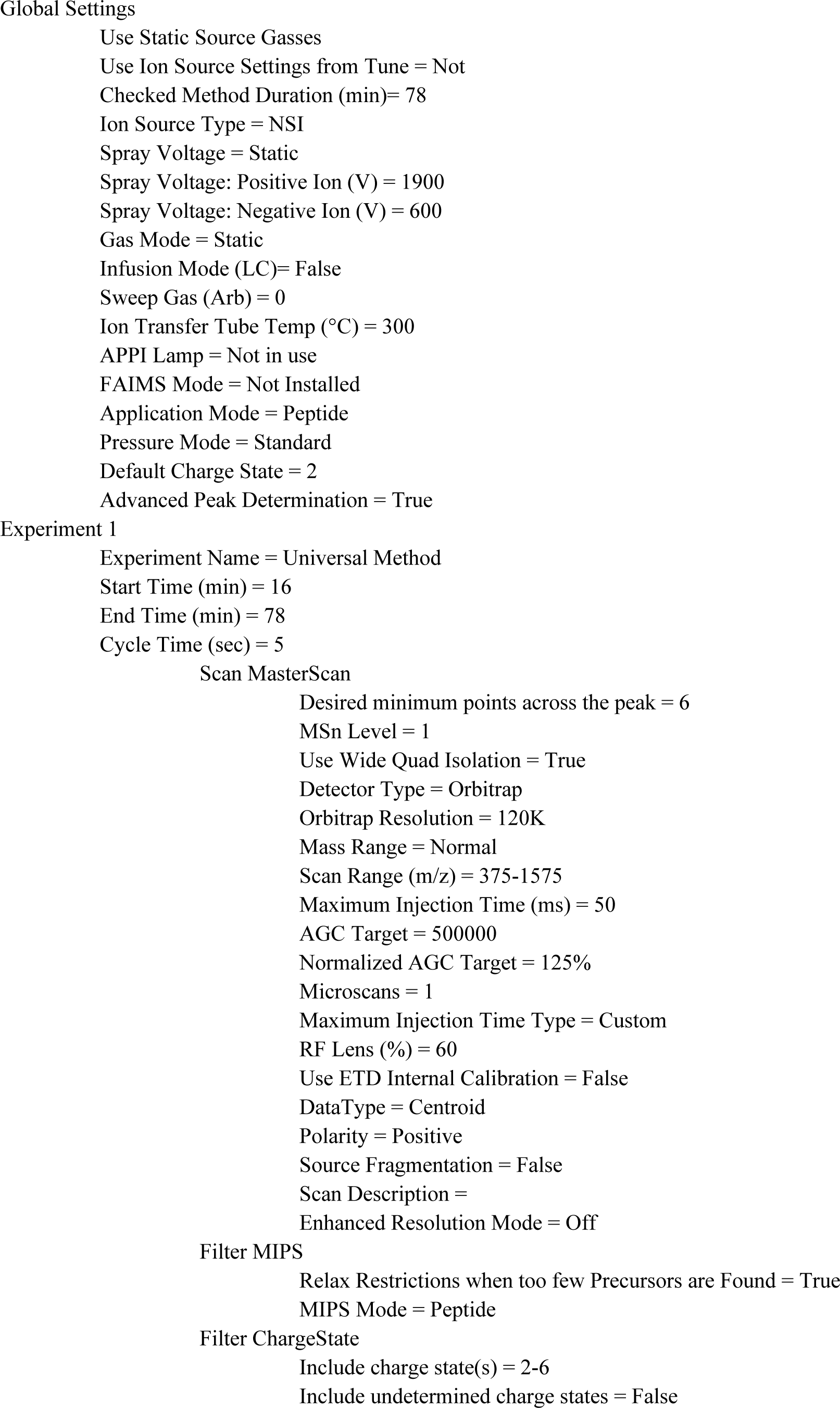
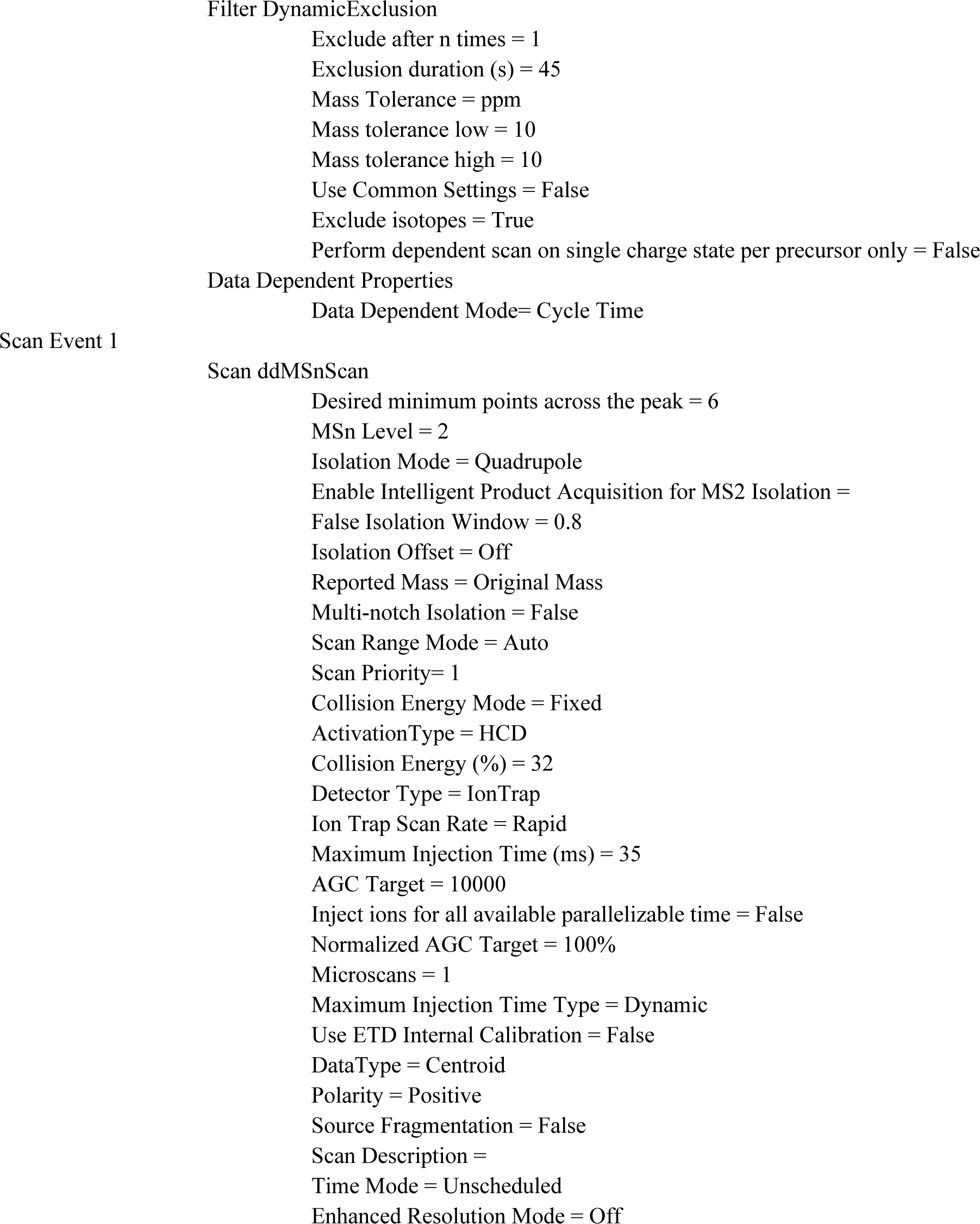
Orbitrap Fusion Method Summary.

## Supplementary Figures

**Figure S1.**
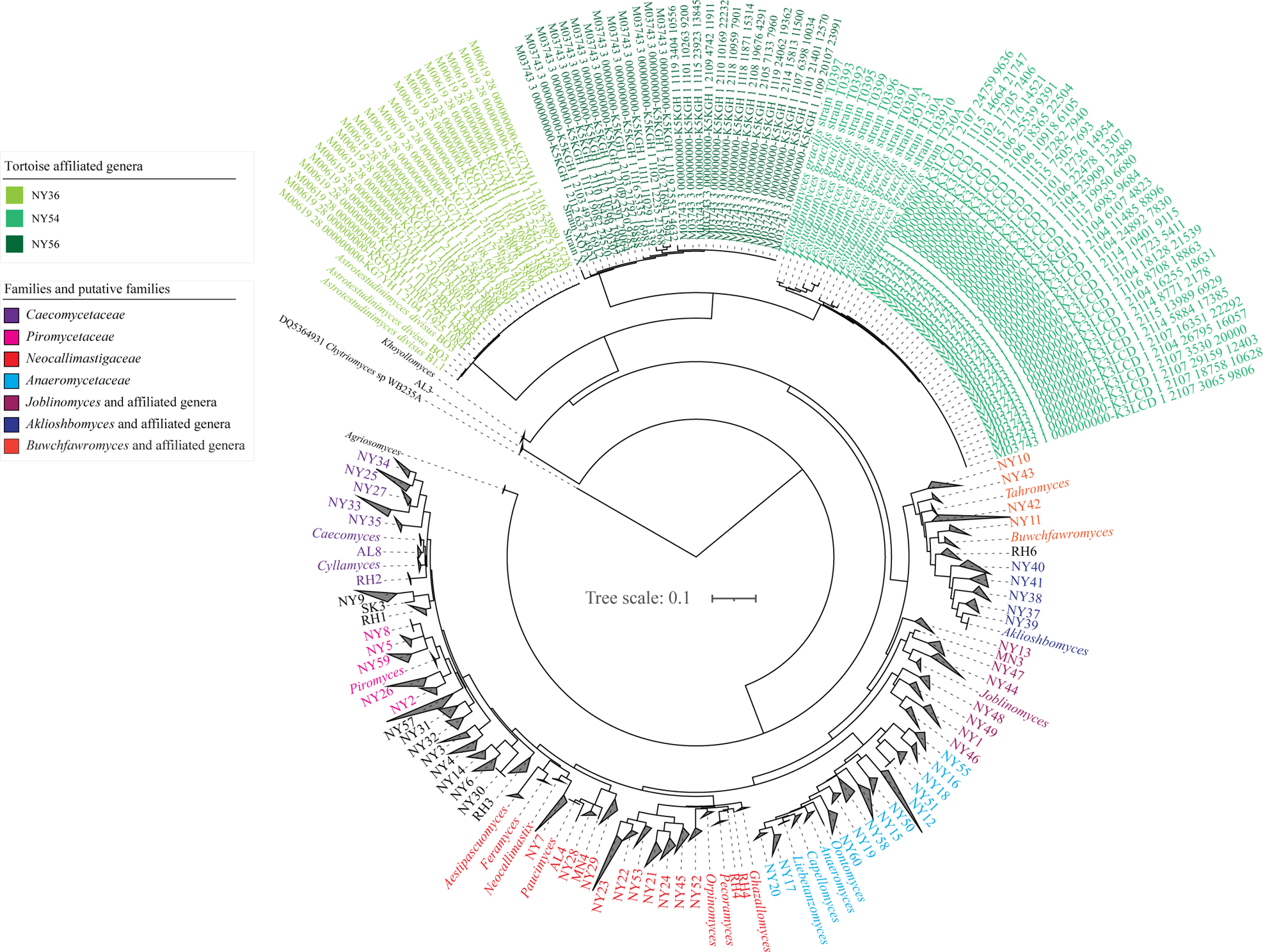
Maximum likelihood phylogenetic tree in Figure 1D with the wedges of the three tortoise affiliated genera expanded and including sequences from the current culture-independent study. All other genera are shown as collapsed wedges and names are color coded by genus as shown in the figure legend.

**Figure S2.**
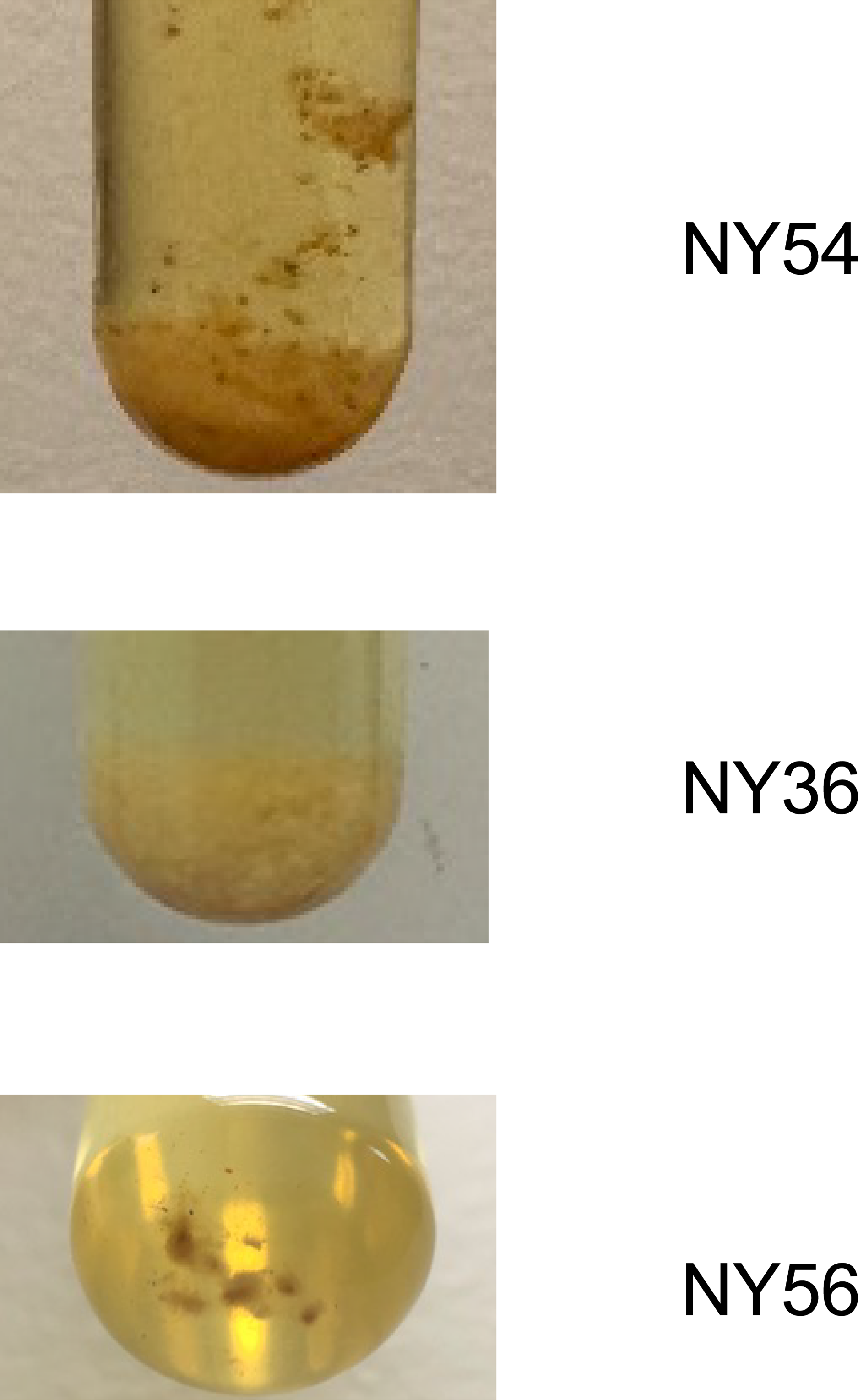
Isolates belonging to the three putative genera growing in liquid RFC media. Isolate names and genus are indicated on the right of each tube.

**Figure S3.**
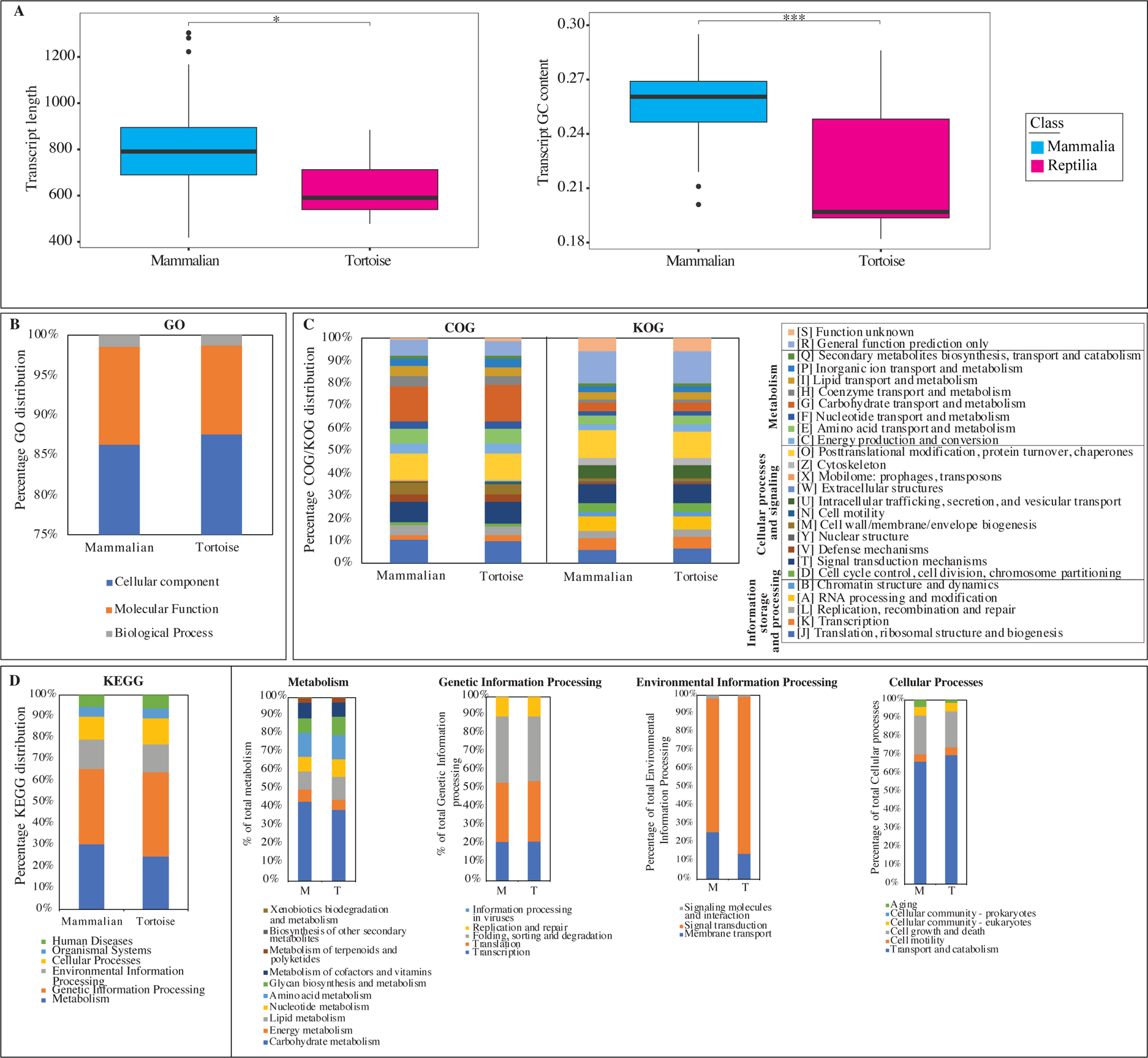
Comparative general features and gene content analysis of the 7 tortoise sourced transcriptomes generated in this study (pink), versus the 52 mammalian sourced transcriptomes generated previously (cyan) (8, 11, 13–16, 24, 25). (A) Distribution of transcript length (left) and GC content (right). Results of two-tailed ANOVA for pairwise comparison are shown on top. (B-C) Gene content comparison between mammalian sourced (left stacked columns) and tortoise sourced (right stacked columns) transcriptomes using GO (B), COG/KOG (C), and KEGG (D) classification. KEGG classification is further broken down into the four main categories: Metabolism, Genetic Information Processing, Environmental Information Processing, and Cellular Processes.

**Figure S4.**
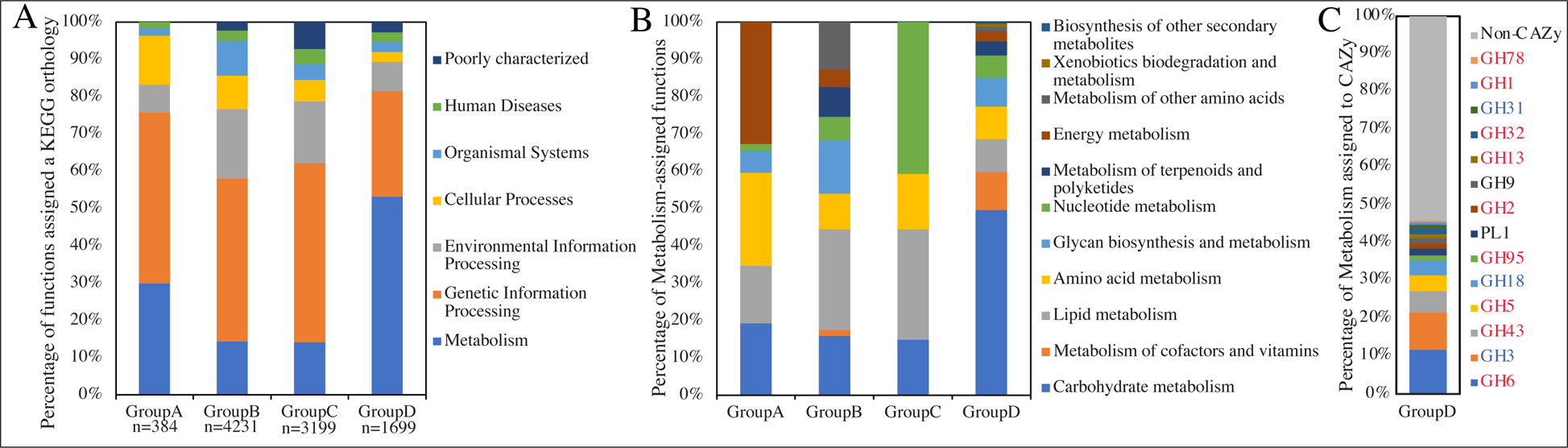
Functional classification of MCL obtained clusters. Four groups of clusters are compared: GroupA: distinct transcripts that are present in both tortoise isolates but absent from mammalian affiliated AGF isolates, n=384 functional clusters; GroupB: distinct transcripts that are present in NY36 but not NY54 or the mammalian affiliated AGF isolates transcriptomes, n=4231 functional clusters; GroupC: distinct transcripts that are present in NY54 but not NY36 or the mammalian affiliated AGF isolates transcriptomes, n=3199 functional clusters; GroupD: distinct transcripts that are present in mammalian affiliated AGF isolates but absent from both tortoise affiliated AGF isolates, n=1699 functional clusters. (A) KEGG classification of clusters in the 4 groups. (B) Zoom in on clusters assigned a KEGG metabolism function for each of the four groups of clusters in A. (C) CAZyome classification of clusters assigned a KEGG carbohydrate metabolism in GroupD clusters. CAZy families previously shown to be completely acquired via HGT are in red text, while families previously shown to be partly acquired via HGT are in blue text (13).

**Figure S5.**
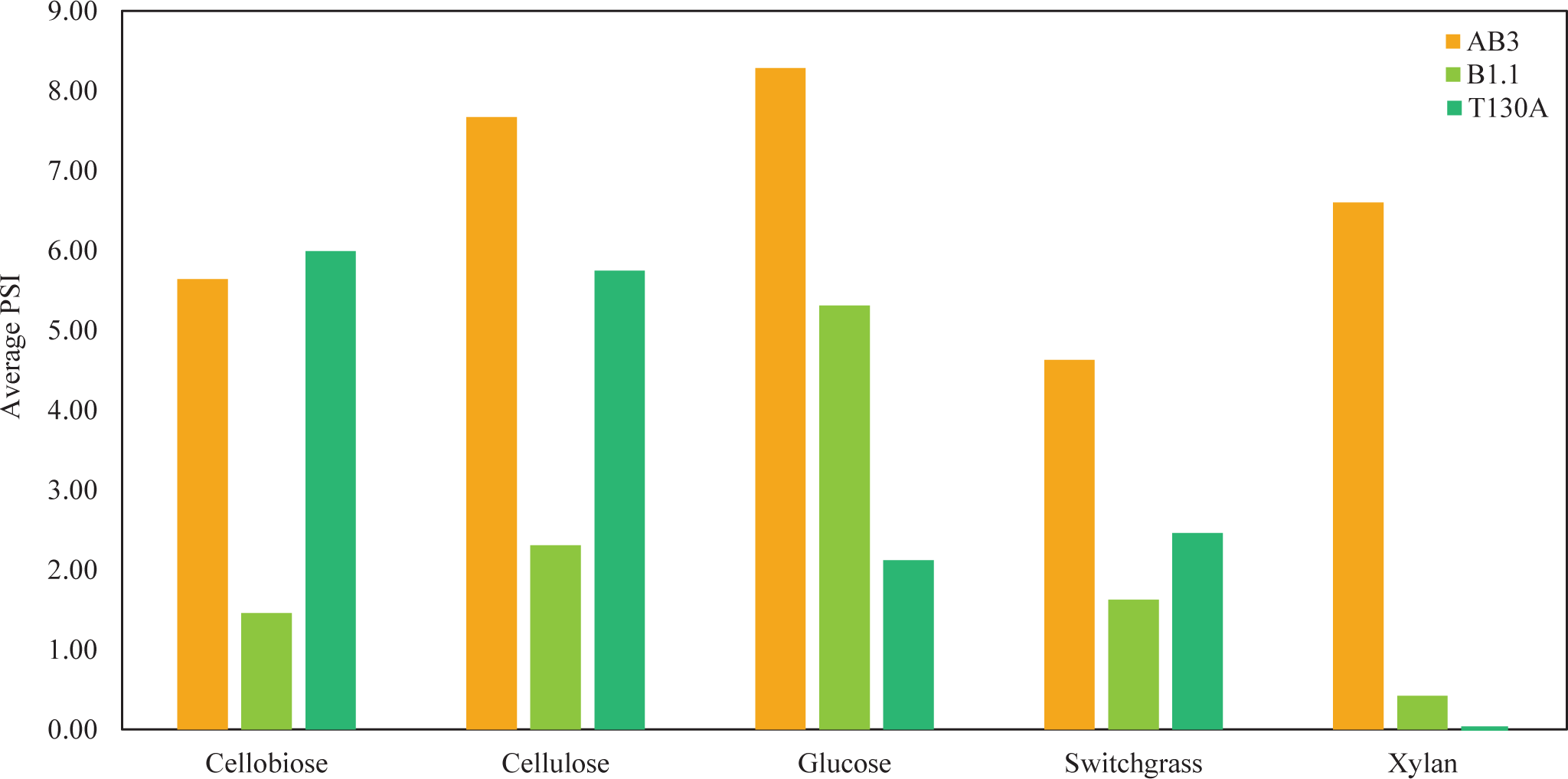
Substrate utilization preferences of two representative isolates of each of the tortoise affiliated genera (NY36 strain B1.1, and NY54 strain T130A) in comparison to an *Orpinomyces joyonii* strain isolated from an American bison (strain AB3). Average gas pressure in PSI (as proxy for growth) from 4 independent growth experiments is shown on the Y-axis, while the carbon source used for growth is shown on the X-axis.

**Figure S6.**
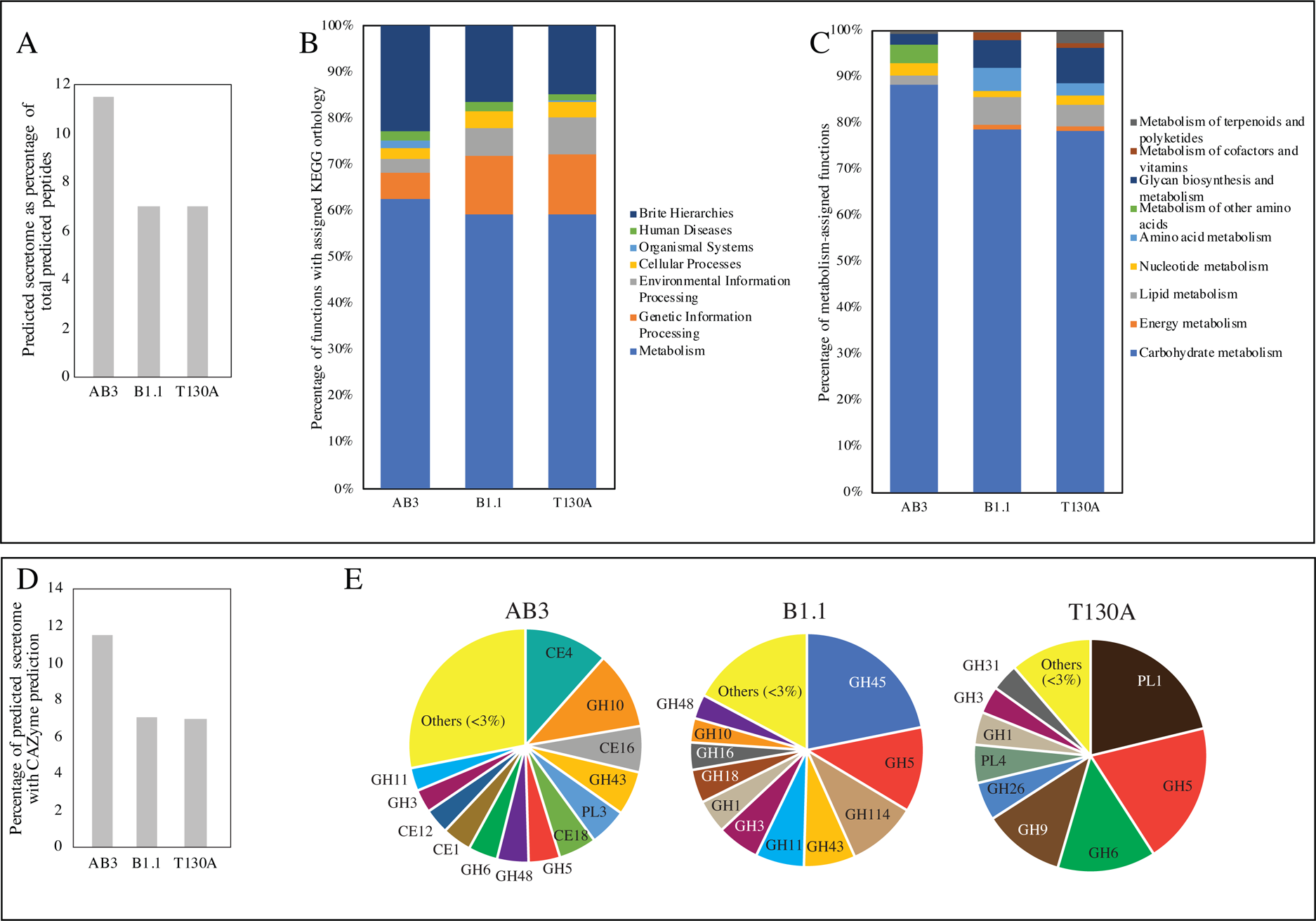
Comparative secretome analysis. The predicted secretome (transcriptome predicted peptides destined to the extracellular milieu as predicted by DeepLoc) of a mammalian AGF isolate, *Orpinomyces joyonii* strain AB3, to these of the tortoise isolates B1.1, and T130A (each representing one of the AGF affiliated genera NY36, and NY54, respectively). (A) Predicted secretome as a percentage of total predicted peptides. (B) Functional classification of the predicted secretome in the three strains. (C) Zoom in on the predicted secretome in the three strains assigned a KEGG metabolism function. (D) Percentage of the predicted secretome in each strain with a CAZyme family prediction. (E) CAZyome composition of of the predicted secretome in each strain. All CAZYme families making up <3% of the total secretome CAZyome are grouped in “others” category.

**Figure S7.**
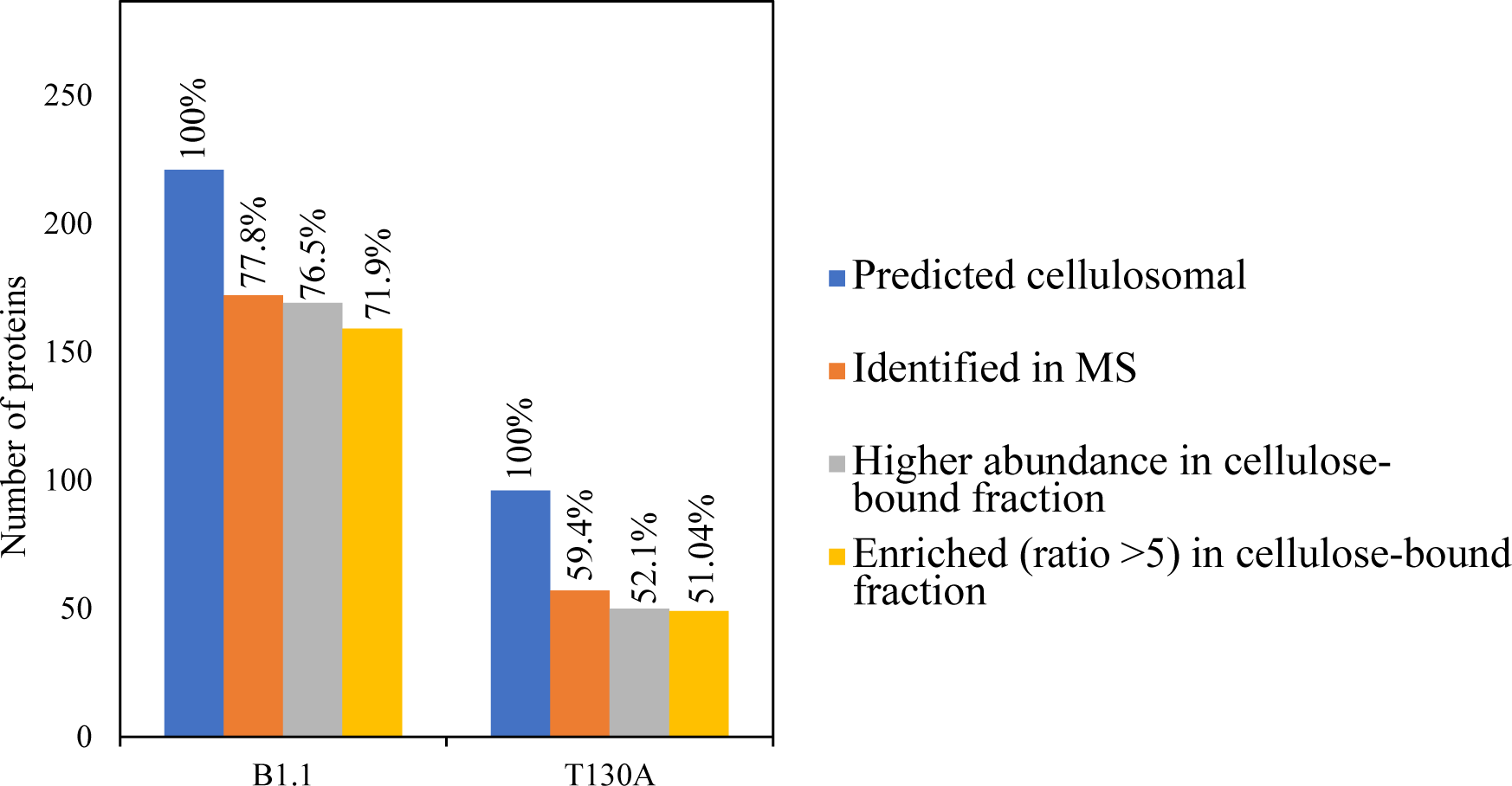
Number of peptides predicted to be cellulosomal in the two tortoise affiliated strains. The blue bars depict the total number of peptides predicted to be cellulosomal from the transcriptomic analysis, the orange bars depict the number of cellulosomal proteins identified in the MS dataset (with the percentage of total proteins shown in top), the grey bars depict the number of proteins found to be with higher abundance in the cellulose bound fraction (ratio of cellulose-bound: biomass intensity >1), and the yellow bars depict the number of peptides found to be enricged in the cellulose bound fraction (ratio of cellulose-bound: biomass intensity >5).

